# A calcium-fingerprint atlas infers the signaling modes of complex elicitors and reveals a salt-like response to a humic biostimulant

**DOI:** 10.64898/2026.10.06.756665

**Authors:** Milan Župunski, Rik Brugman, Martin Zauser, Konstantinos Lampou, Rainer Waadt, Marjorie Guichard, Melanie Krebs, Karin Schumacher, Richard Lamar, Jürgen Pahle, Guido Grossmann

**Author notes:** Co-first author (Milan Župunski, Rik Brugman, Martin Zauser).

## Abstract

A central challenge in plant signaling is to infer, from an observed response, the mechanism that produced it, including perception and signaling components involved. Cytosolic calcium (Ca²⁺) is well suited to this reverse-engineering problem: it is a universal early messenger whose spatiotemporal pattern, the Ca²⁺ signature, is shaped by the specific channels and receptors a stimulus activates. We hypothesize that multiparametric analysis of Ca²⁺ signals can identify the main signaling pathways activated by complex or poorly defined elicitors such as biostimulants. Here, we establish a quantitative framework to resolve Ca²⁺ dynamics in roots of *Arabidopsis* seedlings by transforming spatiotemporal Ca²⁺ transients into multidimensional signatures. Using a normalization and feature extraction approach (Crestline), we define descriptors capturing the onset time, location along the root axis, duration, and propagation of Ca²⁺ signals tip- or shootward, enabling robust comparison across stimuli. Integration of these descriptors reveals stimulus-specific “calcium fingerprints” that allow systematic classification of responses to biotic and abiotic cues within a structured signaling space. To test whether projecting an uncharacterized input into this space can predict its signaling mode, we analyzed a complex humic biostimulant. Its response fell within the fast-onset region of signaling space shared by DAMP and salt responses, clearly apart from MAMP responses, and was altered in *moca1*, a mutant impaired in salt sensing. Together, these findings establish Ca²⁺ dynamics as an interpretable encoding layer for environmental information and provide a scalable framework to resolve complex biological inputs.

## Introduction

Calcium ions (Ca^2+^) represent the most versatile second messenger in cellular signaling across eukaryotes (Berridge et al., 2000; Kudla et al., 2018; Luan, 2026; Župunski et al., 2026). Transient increases in cytosolic Ca²⁺ concentration ([Ca^2+^]_cyt_) are triggered by diverse stimuli, including nutrient availability, salt, drought, temperature fluctuations, herbivory, and microbial elicitors (Behera et al., 2017; Toyota et al., 2018; Tian et al., 2019; Fu et al., 2022; Luan, 2026). These signals are actively shaped by channels and transporters at the plasma membrane and organellar membranes, and are decoded by Ca²⁺-binding proteins such as calmodulins, CPK/CDPKs, CBLs, and CIPKs to regulate downstream responses (Liu et al., 2020; Tang et al., 2020). Plant cells store calcium in the apoplast, the vacuole, and the endoplasmic reticulum, and draw on it through distinct channels at each membrane (Stael et al., 2012). At the cell surface, cyclic nucleotide-gated channels (CNGCs) and glutamate receptor-like channels (GLRs) carry much of the calcium that enters the cytosol after a stimulus (Dietrich et al., 2020; Simon et al., 2023). These channels are gated in different ways. Some open directly when a ligand binds, as GLRs do in response to glutamate (Toyota et al., 2018; Green et al., 2021). Others open indirectly, through second messengers or phosphorylation: cyclic nucleotides and calmodulin gate CNGCs (Fischer et al., 2013; Tian et al., 2019), and calcium-dependent protein kinases activate them downstream of other signals (Zhu et al., 2025; Ming et al., 2025). How long a channel stays open, and how quickly pumps and antiporters return calcium to these stores, together set the timing, spread, and duration of the signal. Because different stimuli engage different combinations of these components, the resulting calcium signals should differ in structure, not just in amplitude.

As shown in roots of *Arabidopsis thaliana* (hereafter *Arabidopsis*), Ca²⁺ signals are not uniform but exhibit stimulus-specific spatiotemporal dynamics (Keinath et al., 2015; Waadt et al., 2017; Waadt et al., 2020), indicating that they function as an information-encoding layer linking stimulus perception to downstream responses (Whalley and Knight, 2013; Lenzoni et al., 2018; Liu et al., 2020). Furthermore, differences in the molecular components of stimulus perception and signal transduction can result in reproducible and distinguishable alterations in the spatiotemporal Ca²⁺ signatures (Wan et al., 2019). Conversely, the characteristics of Ca²⁺ signals often share similarities among responses to elicitors from related classes, e.g. biotic (bacteria, fungi, peptides, cell wall polymers) versus abiotic elicitors (salts, nutrients, water stress, temperature shifts) (Knight et al., 1997; McAinsh and Hetherington, 1998; Ranf et al., 2011; Waadt et al., 2017; Wan et al., 2019; Yuan et al., 2020). These class-level similarities reflect shared perception routes. Microbe- and pathogen-associated molecular patterns (MAMPs/PAMPs), such as bacterial flagellin or fungal chitin, are recognized by cell-surface pattern-recognition receptors that relay through receptor kinases to trigger Ca²⁺ influx (Miya et al., 2007; Ranf et al., 2011; Yuan et al., 2020). Damage-associated molecular patterns (DAMPs), such as extracellular adenosine triphosphate (eATP) or wound-released peptides, are perceived by distinct receptors but converge on a similar Ca²⁺-influx machinery (Tanaka et al., 2014; Harris and Mou, 2024). Abiotic cues act more directly, with salt, osmotic and cold stimuli gating Ca²⁺ entry through membrane- and mechanosensing mechanisms rather than dedicated receptors (Yuan et al., 2014; Jiang et al., 2019). The basis for stimulus-specific Ca²⁺ signatures therefore lie mainly in these different perception mechanisms.

We therefore hypothesized that Ca²⁺ signatures can be used to infer the signaling pathways activated by elicitors, including unknown compounds and complex mixtures such as biostimulants. To test this hypothesis, we combined medium-throughput imaging with quantitative spatiotemporal analysis of Ca²⁺ dynamics to establish a calcium response atlas. We used roots of *Arabidopsis* seedlings because they enable standardized, parallelized imaging and are highly amenable to quantitative image analysis. Roots are the primary interface with the soil environment and integrate diverse biotic and abiotic cues, making them a key site for environmental sensing and Ca²⁺-based signal integration. The calcium response atlas (hereafter termed “Ca²⁺tlas”) was built from datasets of known elicitors. This atlas serves as a reference (ground-truth), enabling multivariate co-clustering analyses to classify responses to novel compounds.

Spatiotemporal dynamics of Ca²⁺ transients, have been measured by high-resolution imaging using genetically encoded Ca²⁺ biosensors, including Yellow Cameleon 3.6, GCaMP3/6, R-GECO1, and NEMO, enabling the spatiotemporal mapping of Ca²⁺ signals in roots at cellular resolution (Krebs et al., 2012; Keinath et al., 2015; Waadt et al., 2017; Liu et al., 2017; Vincent et al., 2017; Li et al., 2023). Kymographs, which compress time-lapse image stacks along a defined root axis, reveal spatial and temporal features of Ca²⁺ transients but are limited by low signal-to-noise ratios and do not readily support quantitative comparison across conditions. Despite advances in imaging, a standardized framework to systematically quantify and compare Ca²⁺ fingerprint dynamics across stimuli is lacking.

To address this, we developed “Crestline”, a signal normalization and feature extraction approach that enhances local Ca²⁺ dynamics and generates “crestplots” that resolve the shape, timing, and propagation of Ca²⁺ signatures across roots. These crestplots define stimulus-specific Ca²⁺ fingerprints, which we quantify using six descriptors – signal onset time (SOT), spatial origin (DfRT), duration (SD), propagation distance (SSS), and tipward (Vt) and shootward (Vs) velocities –providing a multidimensional representation of Ca²⁺ dynamics. Because building a reference atlas requires many roots per stimulus, we applied this framework to medium-throughput, low-resolution recordings rather than to high-resolution imaging of single roots, and all descriptors reported here derive from that dataset. This provides a quantitative basis for comparing Ca²⁺ responses across stimuli. We hypothesized that spatiotemporal Ca²⁺ dynamics define a structured signaling space in which a stimulus is encoded by its position, allowing its dominant signaling class to be inferred. Diverse stimuli, including complex biostimulant mixtures, can then be placed within this Ca²⁺tlas space and compared directly against the reference panel.

## Results

### Amplitude-only quantification obscures stimulus-specific spatial information

To establish a systematic imaging-based analysis workflow of calcium signatures, we first assessed the limitations of amplitude-based [Ca^2+^]_cyt_ quantification in root tissues. In plant roots, exposure to environmental stimuli typically triggers a rapid rise in [Ca^2+^]_cyt_, followed by a resting phase and return to baseline (Fig. 1A). Notably, different elicitors, such as 150 µM eATP and 100 µg mL⁻¹ chitin, produced calcium transients of similar overall amplitude but with clear differences in the timing of the peak response (Fig. 1A). Moreover, amplitude-based readouts were highly sensitive to ROI placement in the spatially heterogeneous root tip. Comparing centerline- and edge-positioned ROIs (5-px width) altered both the apparent response amplitude and the inferred dynamics for the same elicitor (Fig. S1A,D), indicating that amplitude-only measurements can mask underlying spatiotemporal patterns. Together, these observations show that peak-amplitude measurements alone incompletely capture the spatial and temporal complexity of cytosolic calcium responses and can be strongly influenced by ROI placement. Spatiotemporal analysis is therefore required to robustly distinguish and characterize root calcium signatures at the tissue scale.

**Figure 1.**
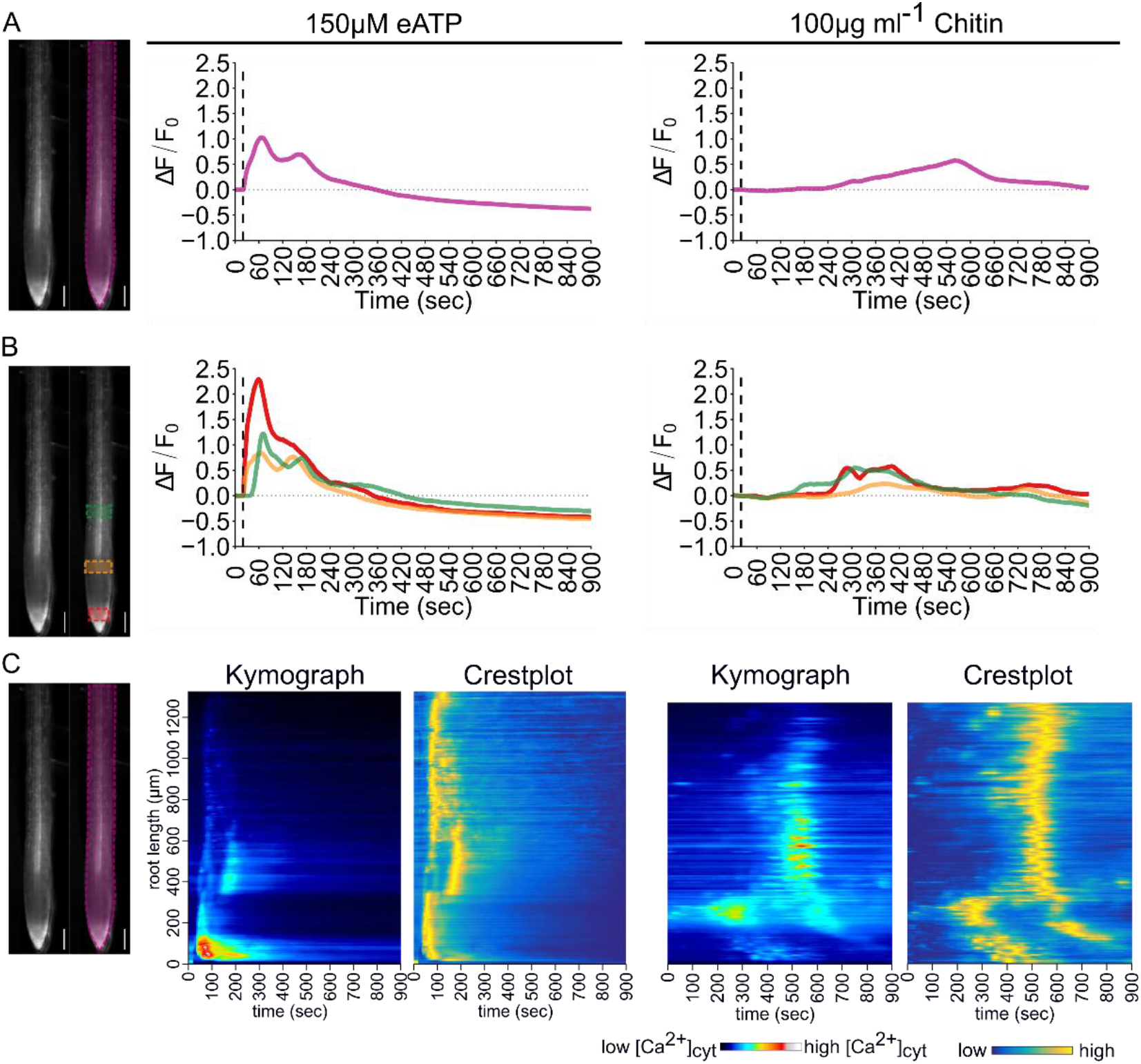
Amplitude-only quantification of R-GECO1 responses misses the spatial dynamics resolved by Crestline. All panels derive from the same two recordings; one root treated with 150 µM eATP (left column) and one with 100 µg mL⁻¹ chitin (right column). A) Conventional ROI-based fluorescence traces (ΔF/F₀) for root tips stimulated with 150 µM eATP or 100 µg mL⁻¹ chitin. The ROI used for intensity extraction is indicated on the root image (scale bar, 50 µm). B) Subdivision of the root into longitudinal zones (color-coded) illustrates that the apparent Ca²⁺ response amplitude depends on ROI position along the root axis, motivating spatially resolved analyses (scale bar, 50 µm). C) Corresponding kymographs (royal LUT) displaying Ca²⁺ signal intensity as a function of time (x-axis) and position along the root (y-axis, root tip at the bottom; 1 pixel column = 1 s), alongside crestplots (wave_tracking LUT) derived from the same recordings, capturing the spatiotemporal trajectory of the Ca²⁺ fingerprint (scale bar, 50 µm). The royal LUT scale spans a fixed display range (min 100 to max 800, arbitrary units) applied identically to all panels, so intensities are directly comparable between them. The wave_tracking LUT spans the full 16-bit output range of the Crestline normalization (0 to 65,535) in all panels. Values are normalized over time at each root position, so they represent relative signal within each position’s time course rather than absolute [Ca²⁺]_cyt_. See Fig. S1 for additional ROI placements and their impact on measured amplitude and inferred spatiotemporal patterns.

When the analysis was shifted from a whole-root perspective to defined developmental zones, spatial differences became even more apparent. In several cases, variation in signal amplitude within a stress class exceeded that observed between stress classes (Fig. 1B), emphasizing the importance of spatial context for interpreting root [Ca^2+^]_cyt_ responses. To capture this information, we generated spatiotemporal maps (kymographs), which enable high-resolution visualization of calcium signaling along the root axis (Fig. 1C). At this resolution, different stimuli produced not only distinct calcium transients, but also stimulus-specific spatial patterns. For example, 150 µM eATP triggered an early and strong signal in the meristematic region, whereas the response appeared later in more differentiated tissues. Likewise, 100 µg mL⁻¹ chitin induced a delayed signal in the elongation zone that subsequently spread along the root.

Although kymographs highlighted the spatial specificity of calcium transients, they could still suffer from low signal-to-noise ratios or variability in signal intensities between measurements, among specimen or within individual roots, e.g., due to variation in biosensor expression along the root axis or imaging sensitivity. Local amplitudes in [Ca^2+^]_cyt_ in weakly sensor-expressing root zones that may be highly relevant for local signaling responses can therefore be overshadowed by high expression in other zones. The issue of uneven sensor expression is typically addressed by including a second reference fluorophore in the sensor design, which requires dual excitation resulting in higher photon-input on the sample (Waadt et al., 2017) or, in the case of newer Matryoshka-type designs (Ast et al., 2017; Ejike et al., 2024), simultaneous acquisition using emission beam-splitting and dual-camera detection. We developed Crestline, a localized normalization approach that visualizes local signal dynamics while reducing background variability and without the need of a reference fluorophore. The Crestline algorithm calculates the local minimum and maximum over the entire measurement period for each spatial position along the root axis and uses these to scale the intensity values. This allows the calcium fingerprint to be visualized even in regions where the absolute intensity values are very low. The resulting output, termed a crestplot, accentuates the shape of the calcium fingerprint while suppressing background variation (Fig. 1C). In crestplots, akin to kymographs, the x-axis represents time, while the spatial axis is plotted along the y-axis, with the root tip consistently positioned at bottom left.

### Bulk seedling treatment enables medium-throughput live imaging of root [Ca^2+^]_cyt_ responses

Testing biological hypotheses requires sufficient replication, yet high-resolution [Ca^2+^]_cyt_ imaging is often limited in throughput by a restricted field of view and the loss of temporal resolution associated with sequential XY-position acquisition. To overcome these constraints, we developed BalanceSir, a simple tilting platform that enables rapid and uniform medium exchange while allowing simultaneous imaging of up to 20 roots in a single microscopy session (Fig. S2).

In this workflow, seedlings are grown on a nylon mesh placed on the agar surface, allowing straightforward transfer into an imaging chamber on the day of the experiment (Fig. S2A–E). Growth on meshes with a mesh size of 0.1 mm (see Materials & Methods) resulted in highly parallel root growth, reducing root cross-over and allowing for higher density seeding of *Arabidopsis* seedlings, thus increasing experimental throughput. Once meshes are transferred into the imaging chamber, tilting enables rapid and uniform medium exchange while maintaining imaging geometry (Fig. S2F, Supplemental Movie 1). Returning the platform to its original position restores the imaging geometry and permits synchronized treatment of all roots within the field of view (Supplemental Movies 1 and 2). Importantly, the tilting procedure itself did not induce detectable [Ca^2+^]_cyt_ transients, as shown by the ½ MS mock controls (Figs. 2, S3). Although BalanceSir still requires manual exchange of media and chambers, it substantially increases experimental throughput while preserving the temporal resolution needed for calcium fingerprint analysis.

**Figure 2.**
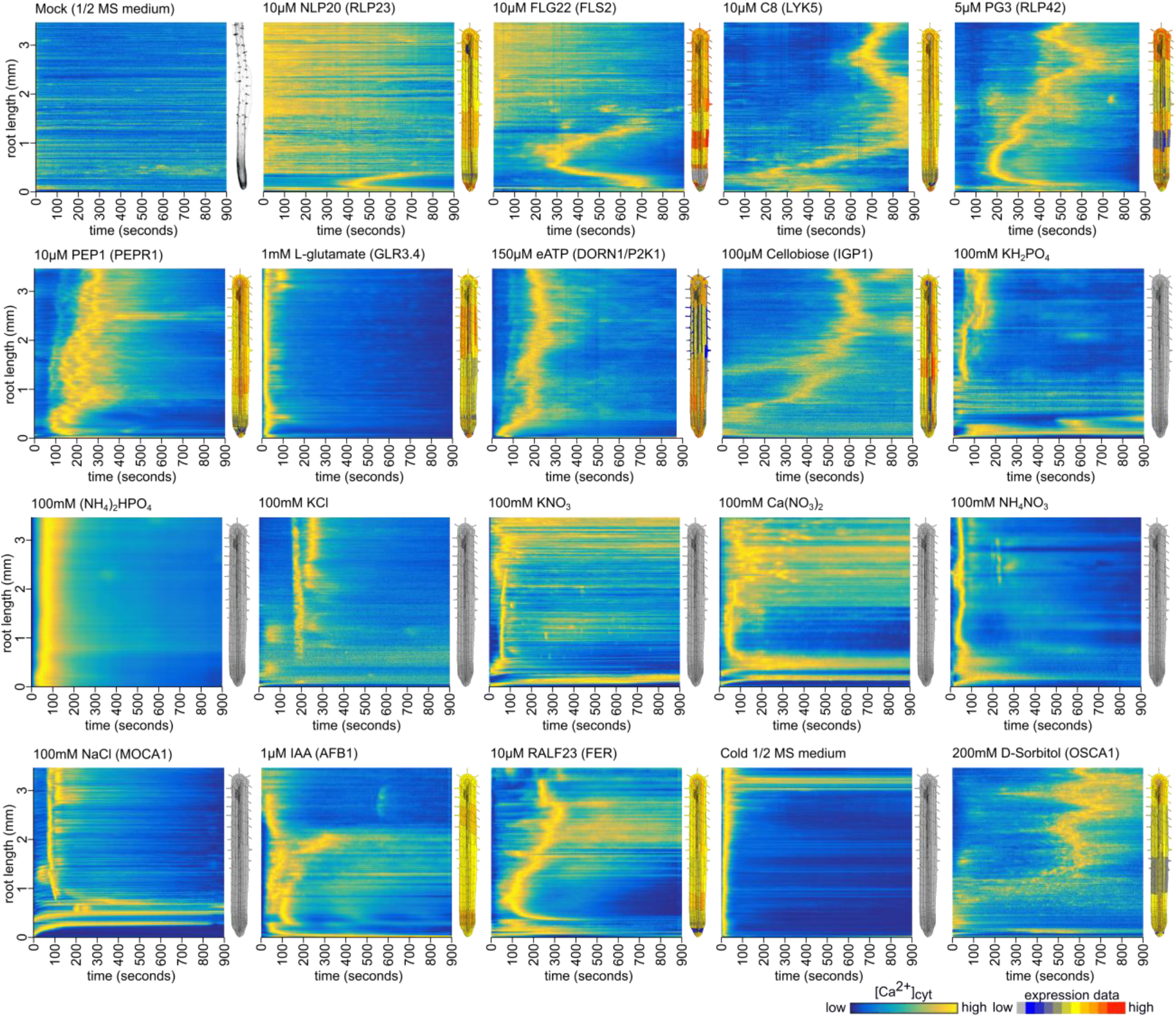
Crestplots reveal distinct, stimulus-specific Ca²⁺ fingerprints across the elicitor panel. Each elicitor and its concentration are indicated above the corresponding panel. The x-axis denotes time (1 pixel column = 1 s), and the y-axis denotes position along the imaged root segment (root tip at the bottom). For elicitors with known perception pathways, the corresponding receptor expression profile is shown next to each crestplot (data from the BAR ePlant resource, https://bar.utoronto.ca/eplant/), with the relevant receptor or sensing component indicated in parentheses; for elicitors with unknown receptors, a grayscale root image is shown instead. A brightfield image of a representative root is shown next to the mock control to indicate root anatomy along the imaged segment. The wave_tracking LUT spans the full 16-bit output range of the Crestline normalization (0 to 65,535) applied identically in all panels; values are normalized over time at each root position, so they represent relative signal within each position’s time course rather than absolute [Ca²⁺]cyt. Expression profiles are shown as relative expression values on the adjacent color scale. They are published reference data (Waese *et al*., 2017), scaled to match the developmental zonation of the imaged roots using the brightfield image as reference; they were not measured in the imaged roots and are shown for exploratory comparison only. Corresponding kymographs are shown in Fig. S3.

### Calcium fingerprint specificity is evident within and across elicitor classes

Building on Crestline normalization, we next asked whether *Arabidopsis* roots respond to different stimuli with distinct [Ca^2+^]_cyt_ patterns, and whether Crestline could resolve these stimulus-specific signatures. To address this, we screened a diverse panel of elicitors representing biotic cues, including microbe-associated molecular patterns (MAMPs) and damage-associated molecular patterns (DAMPs), as well as abiotic cues such as nutrients, cold, salinity, and osmotic stress. This generated a library of spatiotemporal Ca²⁺ responses, hereafter referred to as calcium fingerprints (Fig.2). These fingerprints represent reproducible spatiotemporal patterns that can be formally defined as quantitative multi-parameter descriptors of [Ca^2+^]_cyt_ dynamics.

Crestplots revealed clear stimulus-specific patterns of transient [Ca^2+^]_cyt_ release. For example, the MAMPs NLP20 and FLG22 produced fingerprints that were largely confined to specific root regions, namely the root tip meristematic zone and the elongation zone, respectively, consistent with previously published, comparable responses (Wan et al., 2019). By contrast, the fungal MAMPs C8 and BcPG3 elicited fingerprints that originated in the early elongation zone (approximately 300–500 µm from the root tip) and spread further along the root with distinct kinetics. The DAMPs eATP and L-Glu initiated very rapidly and propagated along the root with fast kinetics. In several cases, these spatial [Ca^2+^]_cyt_ signatures corresponded to known receptor expression domains for NLP20, FLG22 and eATP (Winter et al., 2007; Beck et al., 2014; Albert et al., 2015; Matthus et al., 2022), linking perception sites to fingerprint initiation patterns. Most abiotic stimuli also triggered rapid responses, with the notable exception of 200 mM D-sorbitol, which produced a delayed [Ca²⁺]cyt peak approximately 8–12 min after treatment. Because 200 mM D-sorbitol is approximately iso-osmotic with the 100 mM monovalent salts yet produced a delayed and spatially distinct response, the rapid salt-induced fingerprints are not explained by osmotic load alone, indicating that ionic and osmotic stimuli engage distinct perception routes. Treatment with 100 mM NaCl induced an early primary [Ca^2+^]_cyt_ signal in the maturation/differentiation zone, consistent with previous observations (Steinhorst et al., 2022), followed by a rapidly spreading signal in both directions. By comparison, nutrient salts generally produced less visually distinct fingerprints, most often represented by early, rapidly propagating signals.

The diversity of these root [Ca^2+^]_cyt_ signatures suggested that spatiotemporal dynamics carry information about different environmental cues. To establish the robustness of these fingerprints, we assessed reproducibility across biological replicates and stability across stimulus doses. We therefore screened 14–79 roots per elicitor, depending on the treatment. For representative stimuli of four classes, the core spatiotemporal organization of each fingerprint - the timing, direction, and extent of signal propagation - was consistent across replicate roots, while absolute signal intensity varied between individuals (Fig. 3A). For each stimulus, a median crestplot is shown in false colour, whereas replicate responses are displayed in grayscale to emphasize pattern consistency within treatments. We next assessed dose sensitivity across a range of elicitor concentrations (FLG22 and C8 at 1, 10 and 20 µM; L-Glu at 0.1, 1 and 10 mM; Fig. 3B). Higher doses caused only minor changes in the overall fingerprint pattern across the concentrations tested (Fig. 3B). The degree of dose-sensitivity may itself be an elicitor-specific property, and finer concentration steps could reveal windows of dose-dependence for individual stimuli. In some cases, however, clear threshold behavior was observed; for example, C8 failed to elicit detectable Ca²⁺ transients below a 10 µM concentration. Together, these findings indicate that Crestline captures reproducible, stimulus-specific [Ca^2+^]_cyt_ patterns and provides a robust framework for comparing complex root signaling responses.

**Figure 3.**
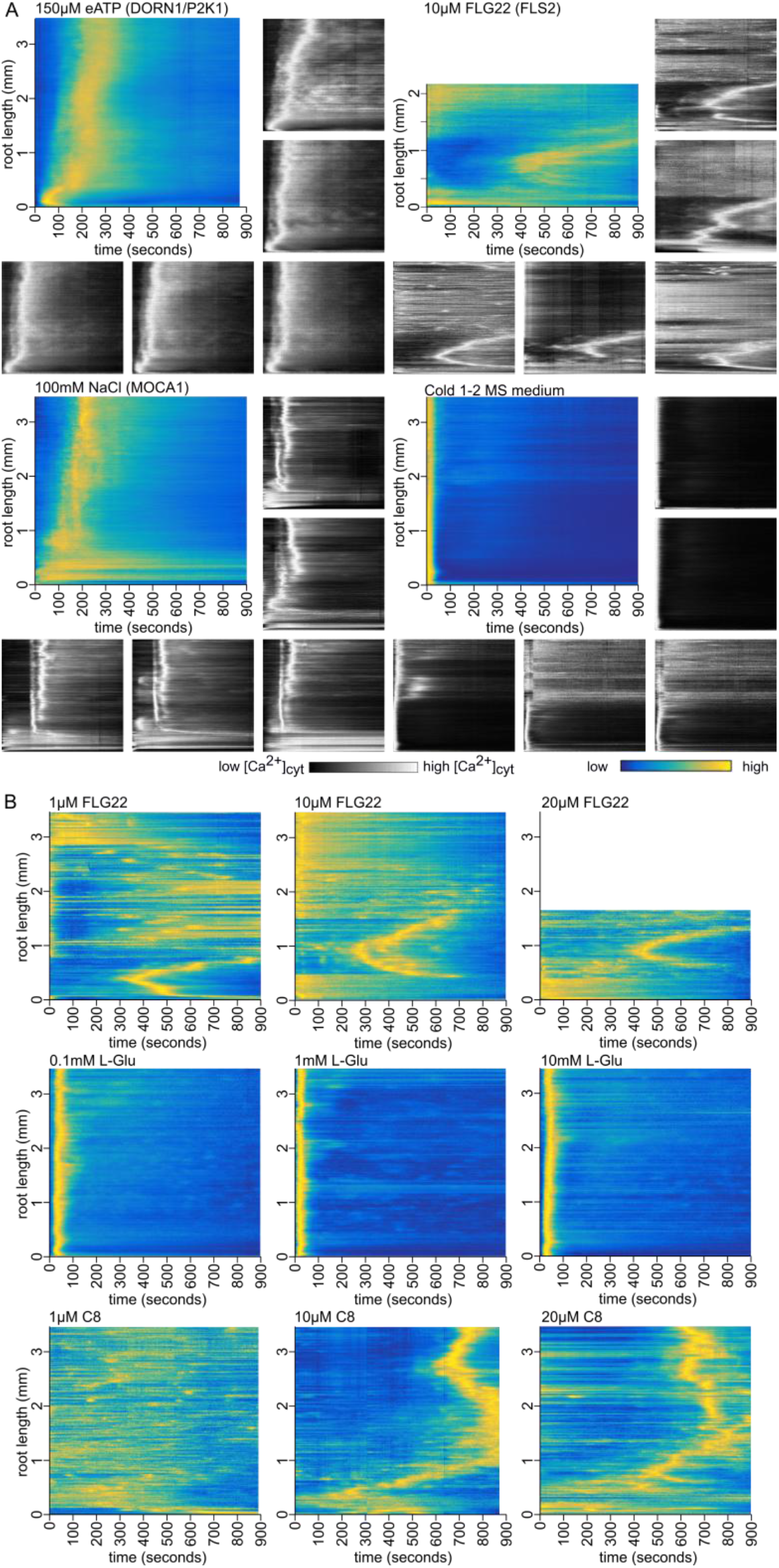
Crestline plots reveal robustness of stimulus-evoked Ca²⁺ fingerprints. A) For four representative stimuli (150 µM eATP, 10 µM FLG22, 100 mM NaCl, cold ½ MS), a median projection of replicate crestplots is shown in the wave_tracking LUT, alongside individual replicate crestplots in grayscale, illustrating the reproducibility of the characteristic Ca²⁺ fingerprint pattern across roots. B) Elicitor-concentration dependence of Ca²⁺ fingerprints. Representative crestplots acquired at different elicitor concentrations show that the core spatiotemporal organization of the Ca²⁺ response is maintained across the concentrations tested. In (A) and (B), the x-axis denotes time (1 pixel column = 1 s), and the y-axis denotes position along the root (root tip at the bottom). The wave_tracking LUT spans the full 16-bit output range of the Crestline normalization (0 to 65,535), applied identically in all panels; values are normalized over time at each root position, so they represent relative signal within each position’s time course rather than absolute [Ca²⁺]_cyt_.

### Quantitative parameterization reveals recurring [Ca^2+^]_cyt_ fingerprint archetypes

To translate spatiotemporal specificity of Ca^2+^ signaling into a quantitative framework and enable inferential comparisons between elicited responses, we next parameterized [Ca^2+^]_cyt_ fingerprints into measurable feature vectors.

We therefore established a pipeline in which multiple aspects of each calcium fingerprint were measured (Fig. 4). Because crestplots capture distinct spatiotemporal features, we defined six quantitative descriptors: (A) the time from stimulus application to first signal detection (SOT); (B) the position along the root at which the first signal appears, (DfRT); (C) the duration of the initial signal (SD); (D) the distance the signal travels from its point of origin (SSS); and (E, F) the velocities of signal propagation through the tissue (Vt and Vs), tipward and shootward, respectively. By simultaneously capturing onset, location, duration, spatial spread, and bidirectional propagation velocities, these six descriptors provide a multidimensional view of calcium fingerprint dynamics, enabling robust discrimination of stimulus-specific fingerprints. To our knowledge, previous studies have generally focused only on a subset of these features, most commonly signature velocity (Choi et al., 2014; Toyota et al., 2018; Bellandi et al., 2022; Steinhorst et al., 2022) or signal delay time (Resentini et al., 2021). Together, this formalizes [Ca^2+^]_cyt_ fingerprint dynamics as quantitative feature vectors that define a structured signaling space for classification and inference.

**Figure 4.**
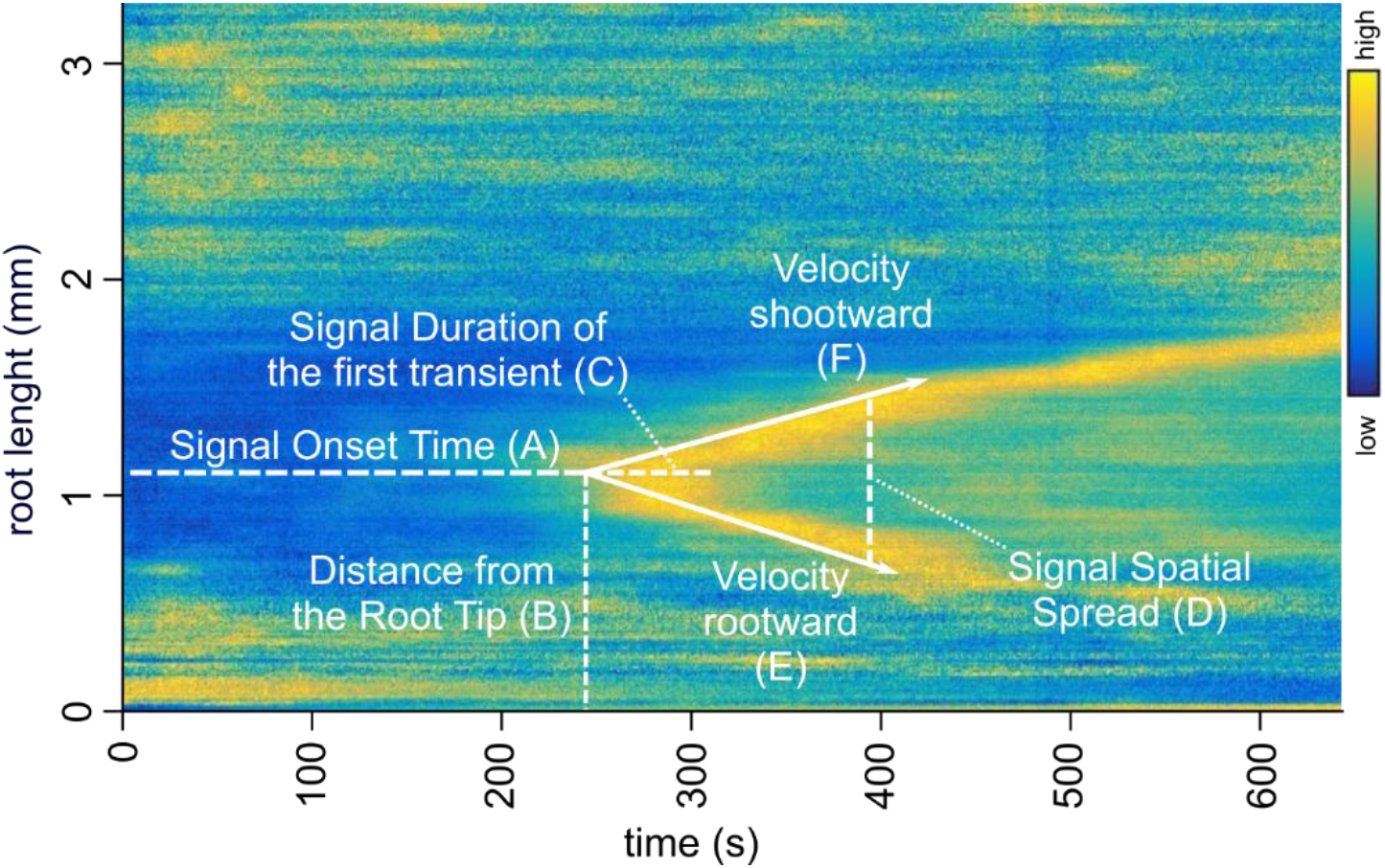
Crestplot parameterization enables quantitative extraction of Ca²⁺ fingerprints. A representative crestplot for 10 µM FLG22 is shown with the spatiotemporal features quantified by the analysis pipeline: (A) signal onset time (delay from stimulus application to first detectable signal), (B) distance from the root tip at first detection, (C) duration of the initial signal, (D) spatial spread (propagation distance from the point of origin), and (E,F) propagation velocities toward the root tip (Vt) and toward the shoot (Vs). The x-axis denotes time (1 pixel column = 1 s), and the y-axis denotes position along the root (root tip at the bottom). The wave_tracking LUT spans the full 16-bit output range of the Crestline normalization (0 to 65,535); values are normalized over time at each root position, so they represent relative signal within each position’s time course rather than absolute [Ca²⁺]cyt.

Quantitative values for all six parameters are shown in Fig. S4. When treatments were arranged according to stimulus class, signal onset time differed strongly across the elicitor panel (Kruskal–Wallis, H = 481.0, df = 18, p = 1.0 × 10⁻⁹⁰), with MAMPs typically inducing delayed responses (medians 137–398 s) compared to most abiotic stimuli (10–80 s), whereas the DAMPs eATP and L-Glu responded almost immediately (medians 4 and 2.5 s) (Fig. S4A). Among the abiotic cues, D-sorbitol was notable for its markedly delayed onset (60–320 s). The site of first detection was also stimulus-dependent (Fig. S4B): some MAMPs and DAMPs initiated close to the root tip (NLP20 and ATP, medians <500 µm), whereas cellobiose and D-sorbitol initiated furthest shootward (medians ∼4,000 µm). MAMPs generally elicited longer-lasting responses (medians ∼60–68 s for FLG22, NLP20 and C8) than the ionotropically perceived DAMP L-glutamate (median 12.5 s), roughly a five-fold difference (Fig. S4C). Analysis of spatial spread further showed that most signals propagated some distance along the root, whereas NLP20 and FLG22 remained comparatively confined to the root tip and early elongation zone, respectively (Fig. S4D). This specificity was also reflected in propagation velocities: NLP20 and FLG22 travelled slowly through the tissue (1–2.5 µm s⁻¹; Fig. S4E,F), whereas nutrient salts produced rapid signatures spreading throughout the root, with tipward velocities ranging from 18.4 to 85.1 µm s⁻¹ and shootward velocities from 63.3 to 96.5 µm s⁻¹.

Together, these analyses revealed two major recurring [Ca^2+^]_cyt_ fingerprint classes across the elicitor panel: (i) slow, spatially confined, and longer-lasting signatures, exemplified by NLP20 and FLG22, and (ii) rapid, long-range propagating signatures, exemplified by nutrient salts and several abiotic cues.

### Ca²⁺ fingerprints form a structured space that separates stimulus classes

Next, we integrated the quantitative descriptors into a multivariate framework that positions elicitors within a shared fingerprint space and enables inference for unknown inputs, thereby testing whether [Ca^2+^]_cyt_ dynamics have predictive power to identify the molecular signaling pathways predominantly engaged by environmental stimuli. Our analysis of elicitor-induced responses in a multivariate space revealed a clustering of broad stress classes, with MAMPs clearly separated from abiotic stimuli and DAMP-associated responses positioned primarily between the two classes (Fig. 5A). This separation indicates that Ca²⁺ fingerprints are sufficiently distinct to enable discrimination between stimulus classes in descriptor space. The intermediate placement of DAMP responses is consistent with the dual nature of DAMPs, which can be released under both biotic and abiotic perturbations. Together, such spatial organization indicates that [Ca^2+^]_cyt_ fingerprints occupy a structured signaling space that enables systematic classification of stimuli based on coordinated descriptor variation. Thus, a stimulus class is defined not by any single parameter, but by a specific combination of parameters within this multidimensional [Ca^2+^]_cyt_ fingerprint space. Examination of descriptor coupling further showed that, across many treatments, signal onset time was negatively correlated with signal duration, and signal duration was negatively correlated with both velocity parameters, with the notable exception of D-sorbitol (Fig. 5B). This descriptor coupling shows that the separation arises from a distributed, multi-parameter structure that amplitude alone cannot capture. To assess whether the Crestline normalization and fingerprint quantification framework depended on the biosensor used, we performed additional experiments with *35S::GCaMP3*, confirming that the qualitative robustness of calcium fingerprints was not specific to *UBQ10::R-GECO1* (Fig. S5A-C).

**Figure 5.**
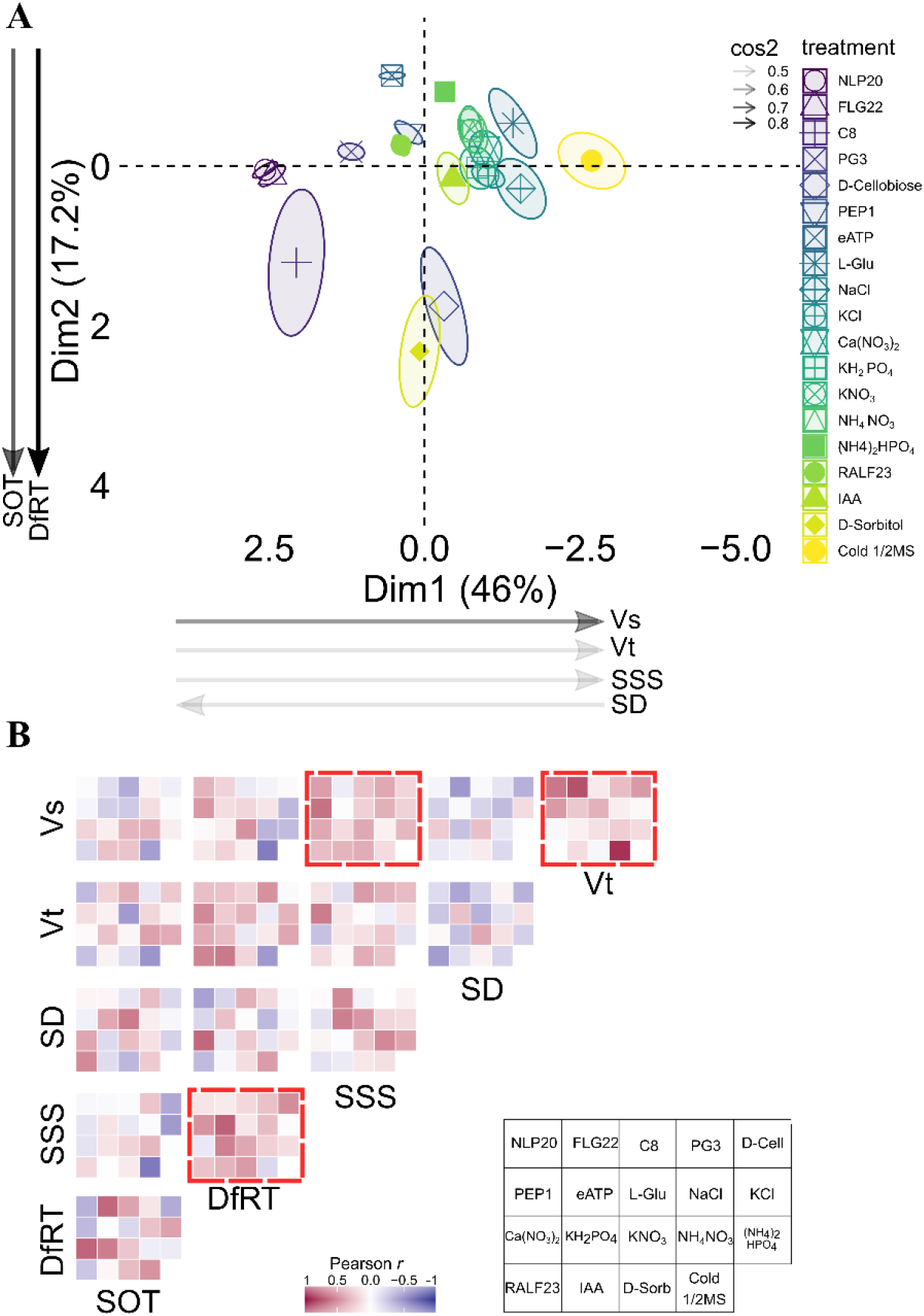
Multivariate structure of Ca²⁺ fingerprint phenotypes elicited by the reference panel of defined stimuli. A) Principal component analysis (PCA) of standardized spatiotemporal Ca²⁺ descriptors extracted from *UBQ10::R-GECO1* imaging along the *Arabidopsis* root tip in response to the defined elicitors (MAMPs/DAMPs, signaling ligands, nutrients/salts, and abiotic stresses). Ellipses show confidence intervals (95%) around the group centroids (means) for each treatment. Vectors indicate the direction of increasing values for the six descriptors used in the embedding: shoot-ward velocity (Vs), tip-ward velocity (Vt), signal spread size (SSS, i.e., propagation length), signal duration (SD), distance from root tip at first detection (DfRT), and signal onset time (SOT). Cos2 denotes contribution of each vector to data spread in multivariate space. Percent variance explained by each dimension is indicated on the axes. B) Within-treatment coupling of Ca²⁺ fingerprint descriptors shown as treatment-specific correlation mosaics (Pearson’s r). Each mini-heatmap represents the pairwise correlation matrix among the six quantitative descriptors (time, distance, length, dur, Vt, Vs) computed across replicate roots for a given elicitor; red boxes highlight descriptor pairs whose coupling differs most across treatments.

### Biostimulant fingerprints fall within the fast-onset region of signaling space

The observation that *Arabidopsis* roots respond to diverse known stimuli with distinct calcium fingerprints raised the possibility that this framework could also be used to analyze unknown or complex inputs and predict the predominant signaling pathways they engage. As a test case, we examined the [Ca^2+^]_cyt_ responses to a humic biostimulant, a class of products known to enhance plant growth under field conditions (Lumactud et al., 2022; Ore et al., 2023; Rathor et al., 2023; Zandonadi et al., 2025). Although broad-range growth-promoting effects for these biostimulants are well documented (Lamar et al., 2024; Santos et al., 2025), the underlying signaling mechanisms remain poorly understood.

We therefore tested the [Ca^2+^]_cyt_ responses to BC4, a proprietary Micro Carbon Technology (MCT) liquid biostimulant containing a blend of organic molecules and colloidal minerals (Huma®, Inc.), using *UBQ10::R-GECO1*, with representative crestplots shown in Fig. 6A. BC4 induced a [Ca^2+^]_cyt_ fingerprint originating in the elongation zone, which was characterized by an early onset and a tipward propagation. To evaluate the compatibility of this response with the established elicitor classes in the Ca^2+^tlas, we projected BC4 recordings into a PCA space computed exclusively on the reference elicitor panel (Fig. 6B). Using the first four principal components (87.4% of total variance explained), we quantified the similarity of each BC4 replicate to the pre-defined elicitor classes by fitting class-conditional multivariate normal models (MVN) in the PCA space and computing posterior class membership probabilities.

**Figure 6.**
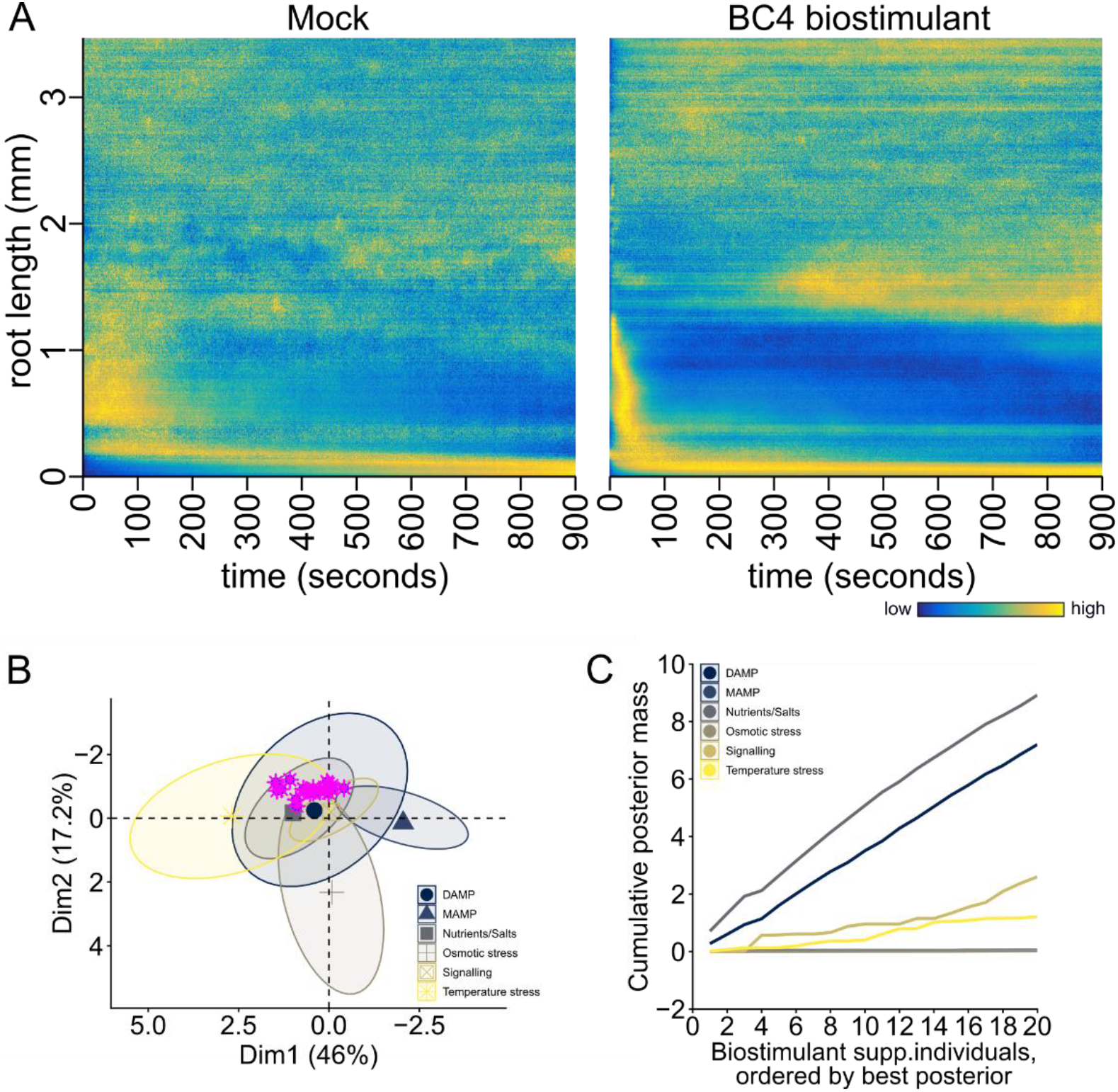
Biostimulant triggers a spatially localized Ca²⁺ response in the *Arabidopsis* root and falls within the fast-onset region of Ca²⁺ fingerprint space. A) Representative crestplots of wild-type Arabidopsis roots expressing *UBQ10::R-GECO1* following mock (½ MS) or biostimulant (BC4, 4 µg C mL⁻¹) treatment. The x-axis denotes time (1 pixel column = 1 s), and the y-axis denotes position along the root (root tip at the bottom). The wave_tracking LUT spans the full 16-bit output range of the Crestline normalization (0 to 65,535), applied identically in all panels; values are normalized over time at each root position, so they represent relative signal within each position’s time course rather than absolute [Ca²⁺]_cyt_. Corresponding representative kymographs are shown in Fig. S6A. B) Principal component analysis (PCA) of Ca²⁺ fingerprint descriptors from the Ca²⁺tlas reference elicitor panel. Reference treatments are grouped by stimulus class (MAMP, DAMP, nutrients/salts, osmotic stress, signaling, temperature stress), and biostimulant replicates are plotted as supplementary individuals, shown in magenta. Ellipses indicate 95% confidence around group centroids. C) Posterior analysis output showing how closely each biostimulant replicate’s Ca²⁺ signature matches each reference stimulus class. Curves summarize the class assignments across all biostimulant replicates under frequency-based class priors: support is split mainly between the nutrients/salts and DAMP classes (see Table S3 for uniform priors).

With class priors proportional to class size, 14 of 20 BC4 replicates were assigned to the nutrients/salts class, 4 to the DAMP class and 2 to the signaling class, with negligible support for the MAMP and osmotic-stress classes (Fig. 6C; Fig. S6B,C). Under uniform priors, support shifted toward the DAMP class (16 of 20 replicates; mean posterior 0.46 versus 0.21 for nutrients/salts), while MAMP and osmotic-stress support remained negligible (Table S3). BC4 support was thus confined mainly to the nutrient/salt and DAMP classes under both priors, with the balance between them depending on the prior. Because a posterior always assigns a sample to some class, we also asked whether BC4 lay within the range of each reference class. By Mahalanobis distance, which does not depend on priors, all 20 BC4 replicates fell within the central 95% of both the nutrient/salt and DAMP classes, nearest to the DAMP centroid, with its nearest reference roots drawn equally from the two classes, whereas BC4 is positioned outside the MAMP and osmotic-stress distributions (Table S3).

To validate the posterior framework, we performed a stratified 5-fold cross-validation on the active reference elicitors using the same class-conditional MVN model. The classifier achieved an overall accuracy of 0.743 and a balanced accuracy of 0.676 (Fig. S6D). Class-wise recall was highest for MAMP (0.903) and remained strong for nutrients/salts (0.780) and signaling (0.797), whereas DAMP showed a lower recall (0.365), consistent with the partial overlap of DAMP Ca²⁺ fingerprints with other stress and signaling classes in this descriptor space. Because stratified cross-validation places replicates of the same elicitor in both training and test sets, we additionally performed leave-one-elicitor-out validation, which tests prediction for stimuli absent from the reference panel. Twelve of 17 evaluable elicitors were assigned to their correct class (root-level accuracy 0.58), and 11 of 17 under uniform priors (0.56; Table S4). All four MAMPs and six of seven nutrient salts were assigned correctly, whereas none of the three DAMPs was, and held-out NaCl was assigned to the DAMP class. The nutrient/salt and DAMP classes therefore overlap in descriptor space and cannot be reliably separated for an unseen stimulus. Together, these analyses place the BC4 response within the fast-onset region of signaling space shared by these two classes, clearly distinct from MAMP-type signaling.

### Genetic perturbation of the biostimulant response

Because the BC4 response fell within the region shared by the DAMP and nutrient/salt classes, with its nearest reference roots drawn equally from both (Table S3), and BC4 contains mineral salts (Table S1), we tested components of salt-induced Ca²⁺ signaling; DAMP perception was not tested here.

The posterior MVN classifier identifies a stimulus class rather than a single pathway; we therefore selected two components implicated in salt/nutrient Ca²⁺ signaling through distinct mechanisms: MOCA1, required for GIPC-sphingolipid-based Na⁺ sensing (Jiang et al., 2019), and CERK1 (Miya et al., 2007; Espinoza et al., 2017) which physically interacts with ANNEXIN 1 (ANN1), a NaCl-induced Ca²⁺-permeable channel, providing a distinct route into salt-triggered Ca²⁺ signaling. We introduced *UBQ10::R-GECO1* into *moca1* and *cerk1-2* backgrounds and established two independent reporter lines per background. To validate reporter functionality in these genetic backgrounds, seedlings were first tested with FLG22, a MOCA1- and CERK1-independent elicitor, and with NaCl, for which altered [Ca²⁺]_cyt_ responses have previously been reported in both mutants (Espinoza et al., 2017; Jiang et al., 2019). All roots of all five lines responded to FLG22, and in the two lines used for the biostimulant experiments (*moca1* #2-1, *cerk1-2* #1-3) onset, duration, spatial spread and velocities of FLG22 signatures did not differ from wild type (Fig. S7C,D; Fig. S8B). NaCl responses differed between lines: *moca1* #1-1 responded with a parameterizable signature in only 11 of 22 roots, which was delayed, slower and less extensive (median onset 40 vs 26 s, shootward velocity 15 vs 120 µm s⁻¹, spread 2.9 vs 4.6 mm), consistent with the nearly abolished NaCl-induced [Ca²⁺]cyt propagation in *moca1* roots (Jiang et al., 2019), whereas *cerk1-2* #1-2 responded almost immediately (median onset 2 s) with a short signature originating closer to the root tip, reminiscent of the hypersensitive response reported for *cerk1* (Espinoza et al., 2017). *moca1* #2-1 and *cerk1-2* #1-3 showed no NaCl fingerprint that could be parameterized (0 of 8 roots each; Fig. S7A,B; Fig. S8A).

To determine whether positioning within the [Ca^2+^]_cyt_ signaling space reflects underlying signaling pathways, we next performed genetic perturbation analysis, quantifying BC4-evoked [Ca^2+^]_cyt_ fingerprint features in one validated reporter line per background (*moca1* #2-1, *cerk1-2* #1-3); both lines responded to the eATP positive control applied at the end of the BC4 recordings (Fig. S9C). In wild type, BC4 treatment produced a characteristic propagating [Ca²⁺]_cyt_ signature in crestplots (Fig. 7A), and in 45% of roots (9 of 20) the initial transient was followed by a second, later signal. In the mutant backgrounds a second signature was not clearly discernible and was therefore not quantified; all descriptors reported here refer to the initial transient. In both mutants the response was shorter, and in *moca1* the signal also spread less. Quantification of spatial spread showed that it differed significantly among genotypes (Kruskal–Wallis, H = 14.2, df = 2, p = 8.2 × 10⁻⁴), with *moca1* signals spreading significantly less than wild type (Dunn’s test, adjusted p = 5.0 × 10⁻⁴) and *cerk1-2* intermediate, not differing significantly from either (Fig. 7B). BC4 treatment also altered signal persistence, with signal duration significantly reduced in both mutant backgrounds relative to wild type (Kruskal–Wallis, H = 18.7, df = 2, p = 8.74 × 10⁻⁵; Fig. 7C; other descriptors, Fig. S9D–G). BC4 signatures in *moca1* initiated slightly but significantly closer to the root tip than in wild type and *cerk1-2* (median distance from the root tip 742 µm vs 856 and 890 µm; Kruskal–Wallis, H = 11.2, df = 2, p = 0.0038; Dunn’s test, adjusted p = 0.028 and 0.0033, respectively), whereas signal onset time and tipward and shootward velocities did not differ among genotypes (Kruskal–Wallis, p = 0.69, 0.16 and 0.48; Fig. S9D-G). Consistent with these effects, PCA of the six descriptors separated *moca1* from wild type, with *cerk1-2* occupying an intermediate position (Fig. 7D). Because Crestline’s min–max normalization removes absolute signal intensity, this separation reflects changes in spatial origin, duration and spread rather than in signal amplitude.

**Figure 7.**
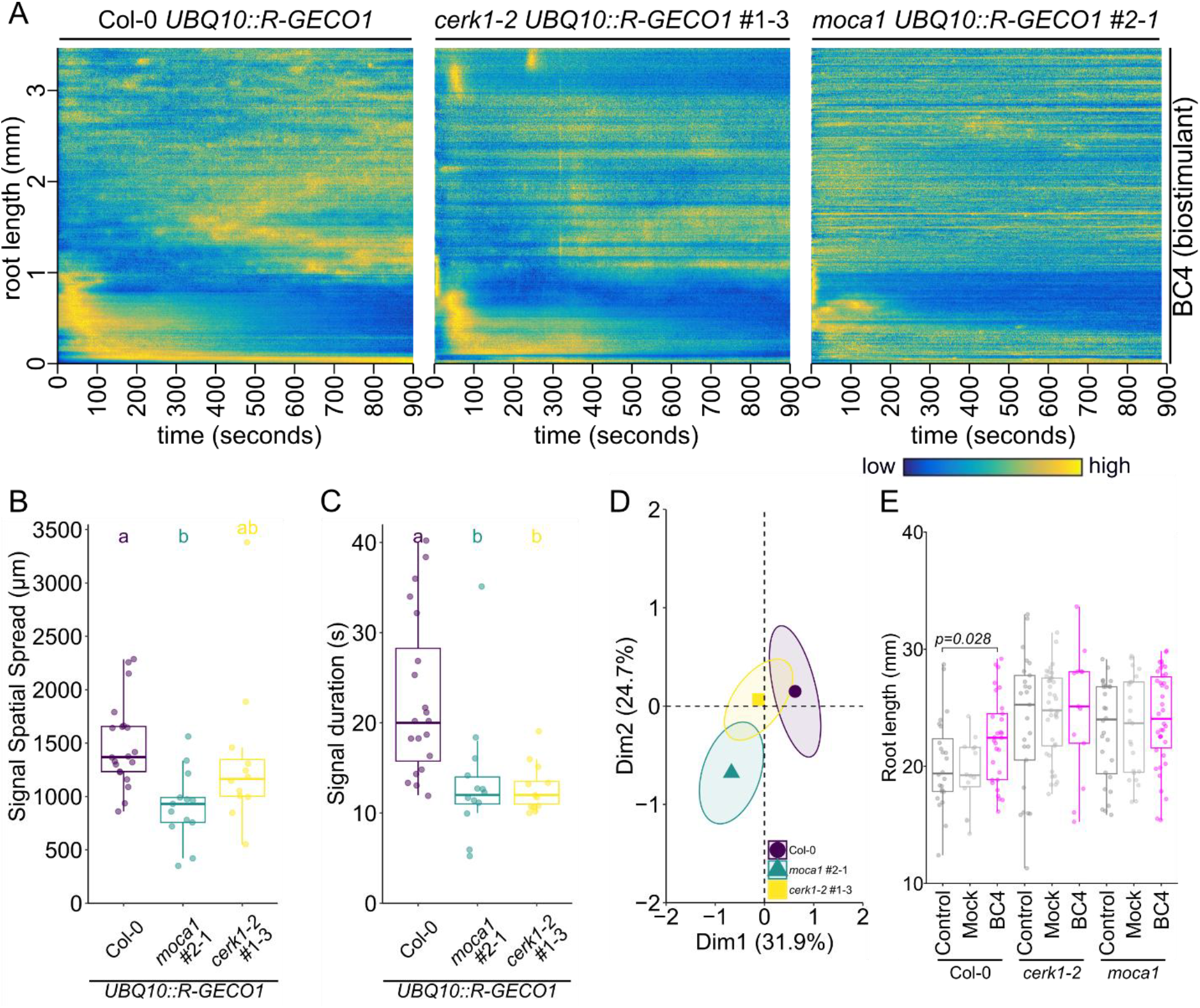
MOCA1 shapes the Ca²⁺ signature evoked by a humic biostimulant in *Arabidopsis* roots. A) Representative crestplots of spatiotemporal Ca²⁺ dynamics in wild type (Col-0), *cerk1-2* (line #1-3) and *moca1* (line #2-1) seedlings expressing *UBQ10::R-GECO1* biosensor, following biostimulant (BC4, 4 µg C mL⁻¹) treatment. The x-axis denotes time (1 pixel column = 1 s), and the y-axis denotes position along the root (root tip at the bottom). The wave_tracking LUT spans the full 16-bit output range of the Crestline normalization (0 to 65,535), applied identically in all panels; values are normalized over time at each root position, so they represent relative signal within each position’s time course rather than absolute [Ca²⁺]_cyt_. Representative kymographs and mock treatment are shown in Fig. S9. B) Spatial spread of the Ca²⁺ signal from its point of origin is genotype-dependent (Kruskal–Wallis, H = 14.2, df = 2, p = 8.2 × 10⁻⁴). Data in (b–d) are from n = 20 (wild type), 12 (*cerk1-2* #1-3) and 13 (*moca1* #2-1) roots. Letters denote post hoc groupings (Dunn’s test with Benjamini–Hochberg correction); all roots responded to BC4. C) Biostimulant (BC4) application differentially affects signal persistence across genotypes, with significant differences in signal duration (Kruskal–Wallis, H = 18.7, df = 2, p = 8.74 × 10⁻⁵). Letters denote post hoc groupings (Dunn’s test with Benjamini–Hochberg correction). D) PCA of Ca²⁺ fingerprint descriptors separate *moca1* from wild type, with *cerk1-2* intermediate. Ellipses represent confidence regions around group centroids. E) The biostimulant (BC4) promotes primary root growth in wild type (parental lines without biosensor: Col-0, *cerk1-2* and *moca1*; linear model, root length ∼ treatment × genotype + batch; estimated marginal means for Col-0, BC4 vs Control +2.8 mm, t₂₂₁ = 2.36, and BC4 vs Mock +3.6 mm, t₂₂₁ = 2.57, both BH-adjusted p = 0.028; Mock vs Control n.s.). The wild-type BC4 effect was confirmed on log-transformed root length (BC4/Control 1.14, BC4/Mock 1.18; both BH-adjusted p = 0.025). The treatment × genotype interaction was not significant (Type II ANOVA, F₄,₂₂₁ = 1.12, p = 0.35; confirmed on log-transformed root length, F₄,₂₂₁ = 1.29, p = 0.27), so no genotype-dependence of the growth response was detected under these conditions. The experimental batch was included as a blocking factor (n = 13–38 roots per condition).

Projected into the Ca²⁺tlas, BC4 responses of both mutants shifted away from the nutrient/salt class (Fig. S11; Table S5). Mean posterior support for this class fell from 0.45 in wild type to 0.38 in *cerk1-2* and 0.30 in *moca1* (Kruskal–Wallis, H = 13.2, df = 2, p = 0.001; *moca1* vs wild type, p = 0.0009), and no *moca1* replicate was assigned to it. Independently of priors, the distance to the nutrient/salt centroid increased in both mutants (H = 20.9, df = 2, p = 3.0 × 10⁻⁵; *moca1* vs wild type, p = 4.6 × 10⁻⁵; *cerk1-2* vs wild type, p = 0.004), and distances to the DAMP, signaling and osmotic-stress centroids increased as well (Table S5). To determine whether the BC4-induced signaling fingerprint was associated with its growth-promoting effects, we quantified root growth in *Arabidopsis* seedlings. BC4 treatment increased primary root length in wild type (+2.8 mm vs control and +3.6 mm vs mock, p = 0.028 for both), whereas no significant growth effect was detected in *cerk1-2* or *moca1*; the treatment × genotype interaction was not significant (Fig. 7E; Fig. S10), so a genotype-dependence of the growth response could not be established under these conditions.

## Discussion

### The Ca²⁺tlas enables differentiation of stimulus-specific calcium signatures

For a long time, the calcium signature hypothesis has been a subject of discussion among researchers (Scrase-Field and Knight, 2003; Plieth, 2005; Kudla et al., 2010; Steinhorst and Kudla, 2013; Plieth, 2016). The work of Whalley & Knight (Whalley and Knight, 2013) experimentally showed that subtle differences in calcium signatures (amplitude, frequency, and number of pulses) enable unique gene expression profiles. However, up to today, there has not been a consensus about how these signatures should be defined ⎼ or even if they exist in the first place ⎼ although the concept has provided a framework aiding in advancing our knowledge and understanding of plant calcium signaling. By quantifying onset time, spatial origin, duration, distance, and propagation velocities in a unified framework, Crestline provides a multi-parametric readout that extends amplitude-only approaches, allowing robust discrimination of stimulus-specific fingerprints and enabling comparative analyses across treatments and genotypes. Our results provide quantitative support for the existence of stimulus-specific Ca²⁺ fingerprints, demonstrating that spatiotemporal Ca²⁺ dynamics are sufficiently distinct to enable systematic discrimination between stimuli. It should be mentioned that previous research also reported spatio-temporal datasets for various elicitors, depicted as kymographs (Krebs et al., 2012; Keinath et al., 2015; Waadt et al., 2017; Waadt et al., 2020) or time-series (Behera et al., 2017; Steinhorst et al., 2022; Fu et al., 2022), although mainly focusing on specific physiological aspects.

Our work demonstrates that *Arabidopsis* roots respond robustly to individual stimuli, producing spatially organized Ca²⁺ signatures even when the entire root is exposed to the elicitor simultaneously (Fig.3A). This argues that specific root tissues, or even cell clusters, serve as sensing niches ⎼ as previously demonstrated for NaCl, Mn, and K (Behera et al., 2017; Steinhorst et al., 2022; Fu et al., 2022). Several elicitors in our panel (MAMPs and DAMPs) are perceived in defined primary-response regions along the root axis. However, perception in that region does not necessarily restrict the downstream response to that region: local immune perception can trigger signaling that spreads beyond the initial site (Zhou and Zhang, 2020; Dodds et al., 2024). Interestingly, for certain elicitors like NLP20 and FLG22, primary-response regions seem to correlate with the expression profiles of their respective receptors (Fig.2). This is particularly interesting for the flagellin epitope FLG22, as Wyrsch et al. (Wyrsch et al., 2015) reported that all root tissues are capable of perceiving FLG22 and inducing PTI responses if the FLS2 receptor is present. According to our quantitative data, most MAMP and DAMP signatures originated near the root tip or in the elongation zone, whereas those of NaCl, several other salts, IAA, cold, D(+)-cellobiose and D-sorbitol originated further shootward (Fig. S4B). Cellobiose is considered a DAMP signal, primarily associated with biotic stress, as it has been shown to induce the expression of defense-related genes, similarly to other pathogen-associated molecular patterns like chitin (Souza et al., 2017).

A Ca²⁺ signature reflects how a stimulus is perceived, which does not always follow the conventional biotic/abiotic grouping. For example, the DAMP cellobiose clustered with the osmotic stimulus D-sorbitol rather than with other DAMPs (Fig. 5A; Table S4), indicating that fingerprint position tracks perception mechanism more than categorical labels. The robustness and spatial specificity of calcium fingerprints suggest that roots possess defined sensing niches capable of perceiving diverse stimuli, while these signals are subsequently integrated to coordinate localized or systemic responses. This highlights the potential of calcium signatures as functional readouts of early perception. The distinct Ca²⁺ fingerprints likely arise from defined perception sites, linking stimulus detection to the onset of specific signaling responses. Above the response threshold, the fingerprint pattern changed little with dose (Fig. 3B), whereas amplitude-based studies report that [Ca²⁺]_cyt_ increases scale with elicitor concentration for FLG22 (Chi et al., 2021; van Dieren et al., 2024), NaCl, mannitol and H₂O₂ (Steinhorst et al., 2022; van Dieren et al., 2024). This suggests that stimulus identity is encoded in the spatiotemporal pattern, whereas dose may be encoded mainly in amplitude.

### Application of the Ca²⁺tlas to infer signaling modes of unknown compounds

For many stimuli, and especially for complex or uncharacterized inputs such as biostimulants, the signal perception and downstream components involved remain unknown. Identifying them has traditionally relied on genetic screens, which are powerful but slow and require a prior analysis about which genes to test. Because Ca²⁺ signatures are shaped by the specific channels and receptors, they offer a different entry point: if an unknown stimulus produces a signature resembling those of stimuli with known perception pathways, that resemblance can predict the likely pathway, narrowing the genetic search before any mutant is made. After establishing calcium fingerprint specificity, we hypothesized that this dataset could be used as a proxy to accelerate the identification of signaling pathways for previously unknown stimuli. In this context, biostimulant compounds seemed reasonable targets, as many products are known to promote growth under field conditions ⎼ albeit with no clear understanding of the underlying mechanisms. Although the Ca²⁺tlas currently comprises 20 conditions (19 elicitors plus a mock control), it spans key biotic and abiotic plant elicitors (Fig. 2). We utilized this quantitative dataset and, within a multivariate space, the biostimulant was placed in the region of signaling space shared by the nutrient/salt and DAMP classes, with the balance between them depending on class priors, and clearly apart from MAMP responses (Fig. 6B,C). Whether the mineral or the organic fraction of the product drives this response was not resolved here. Nevertheless, the biostimulant response was altered in *moca1* despite its low Na⁺ content, raising the possibility, which remains to be tested, that MOCA1 contributes to ionic sensing beyond its established role in NaCl perception. Because eATP and glutamate are released during many abiotic and biotic stresses, the DAMP-like component of the BC4 fingerprint may reflect secondary danger signaling triggered by the treatment itself.

A Ca²⁺ signature can therefore indicate the class of the dominant perceived input. While studies on humic acid compounds often report improved nutrient uptake in plants (Zanin et al., 2018; Jing et al., 2020; Aranaz et al., 2023), they rarely focus on the ionomic profiles of these products. This highlights the potential of calcium signatures to help characterize complex or incompletely defined stimuli. Having the ability to unveil the putative mode of action for certain stimuli, such as humic acids, would advance our understanding of how these compounds are perceived by plants, and utilized to protect crops and stabilize yield. This suggests that Ca²⁺ fingerprints can serve as functional readouts of early signaling processes, enabling data-driven inference of stimulus class and providing a scalable approach to dissect complex biological inputs. We note that these findings are based on *Arabidopsis* roots under controlled conditions; extension to crop species and field settings remains to be demonstrated, and the framework infers the dominant signaling class rather than establishing causal mechanism. The descriptors also capture only the initial signal of each response; secondary calcium signals, such as those observed in nearly half of wild-type roots treated with BC4, are not represented in the signaling space, and BC4 responses of the mutants were quantified in one reporter line each. Finally, like other single-fluorophore Ca²⁺ indicators, R-GECO1 can be influenced by changes in cytosolic pH, which some treatments, notably ammonium salts, may induce; per-profile normalization does not correct for such effects.

Taken together, we establish a quantitative framework for analyzing spatiotemporal Ca²⁺ dynamics that extends beyond amplitude-based interpretation. By transforming Ca²⁺ signals into multidimensional descriptors, we show that Ca²⁺ responses occupy a structured signaling space in which position predicts the dominant class of perception engaged by a stimulus. This provides a proof of concept that the Ca²⁺tlas-derived class prediction can guide targeted genetic validation, narrowing the search for perception and signaling components before a mutant is made. We emphasize that this represents a single worked example rather than a systematically validated predictive pipeline; broader testing across stimuli and pathways will be required to establish the framework’s general predictive power. More broadly, shortening the path from an observed signaling response to its underlying molecular components could move plant signaling research toward systematic, Ca²⁺tlas-guided discovery across the many uncharacterized stimuli plants encounter under field conditions.

## Material and methods

### Plant materials and Growth conditions for Arabidopsis

*Arabidopsis thaliana* (Col-0) lines expressing calcium biosensors: *UBQ10::R-GECO1* (Keinath et al., 2015) and *35S::GCaMP3* (Vincent et al., 2017) were used. The mutant lines *moca1* (Jiang et al., 2019) and *cerk1-2* (GABI-Kat 096F09; Miya et al., 2007) were used to assess pathway-specific responses. To generate reporter lines in mutant backgrounds, the *UBQ10::R-GECO1* cassette was obtained from the original *UBQ10::R-GECO1-mTurquoise* plasmid (Waadt et al., 2017). The selection cassette was subsequently replaced using In-Fusion cloning (Takara Bio) to introduce a FAST-based selection module containing hygromycin resistance and FAST marker (Stuttmann et al., 2021). The resulting construct was transformed into the *Agrobacterium tumefaciens* strain GV3101 and introduced into *moca1* and *cerk1-2* plants via the floral dip method (Clough and Bent, 1998). Transgenic plants were selected based on FAST seed fluorescence and/or hygromycin resistance, and two homozygous lines for both the reporter and the mutant alleles were used for subsequent calcium imaging experiments.

Seeds were surface-sterilized prior to sowing by immersion in 70% ethanol for 15 seconds, followed by 3 minutes of shaking in 6% sodium hypochlorite (NaOCl). After sterilization, seeds were rinsed five times with Milli-Q H₂O and sown either on a sterile nylon mesh (100 µm pore size; Schwegmann Filtration Technology, details in Fig. S2), or, for high-resolution imaging, in an imaging chamber shaped with 3D-printed three-well comb made from polyethylene terephthalate glycol, in which the agar pad and cover glass bottom form a narrow channel for the root.

The growth medium consisted of ½-strength Murashige & Skoog (MS) minimal organic powder medium (pH adjusted to 5.7 with KOH; SERVA Cat. No. 47515.04), supplemented with 0.1% MES monohydrate (Sigma-Aldrich Cat. No. 69892) and 0.8% Plant Agar (Duchefa Biochemie Cat. No. P1001). Seedlings were grown on agar medium in square plates under long-day conditions (16/8 h day/night cycle) for 7 days prior to imaging, under 150 µE LED light intensity, 22/20 °C (day/night), and 65% relative humidity.

### Chemicals, Buffers, and Elicitors

All chemicals were prepared as described below unless stated otherwise. Stock solutions of 100 mM ATP disodium salt (Sigma-Aldrich Cat. No. A2383), 1 M D(+)-cellobiose (Carl Roth Cat. No. 5840.1), 1 mM FLG22 (sequence QRLSTGSRINSAKDDAAGLQIA, purity >95%, Eurogentec Cat. No. AS-62633), 1 mM of the 20-amino-acid epitope of necrosis- and ethylene-inducing peptide 1 (Nep1)-like proteins (NLP20, sequence AIMYSWYFPKDSPVTGLGHR, purity ≥94%, Genscript), 1 mM BcPG3 (purity ≥95%, sequence TGLTCSGGHGLSIGSVGGKSNN, Genscript), 1 mM PEP1 (sequence ATKVKAKQRGKEKVSSGRPGQHN, purity ≥85%, Genscript), 100 µM RALF23 (sequence ATRRYISYGALRRNTIPCSRRGASYYNCRRGAQANPYSRGCSAITRCRRS, purity ≥95%, Genscript), 1 mM chitooctaose (octa-N-acetyl, short C8, purity ≥95%, Isosep Cat. No. 57/12-0005), were dissolved in Milli-Q H_2_O, aliquoted and stored at −20 °C until needed. Stock solutions of 100 mM L-glutamic acid potassium salt (Sigma Aldrich Cat. No. G1501), 1 M NaCl (Carl Roth Cat. No. 3957.1), 1 M KCl (AppliChem Cat. No. A3582), 1 M KH₂PO₄ (ChemSolute Cat. No. 16481000), 1 M D-sorbitol (Sigma-Aldrich Cat. No. S1876), 1 M KNO_3_ (Acros Organics, Cat. No. 424151000), 1 M Ca(NO_3_)_2_×4H_2_O (Fluka, sigma-Aldrich Cat. No. 21197), 1 M NH_4_NO_3_ (Panreac Applichem Cat. No. 131135.1211), 1 M (NH_4_)_2_HPO_4_ (Sigma-Aldrich Cat. No. 215996), were dissolved in Milli-Q H_2_O and stored at −20 °C until needed. 100 mM 3-indoleacetic acid (IAA, Sigma Aldrich Cat. No. I3750) was dissolved in DMSO and stored at −20 °C until needed. The mock for IAA contained 0.001% DMSO. Before use, the stock was serially diluted in ½ MS medium to the final concentration of 1 µM. The resulting final DMSO concentration was 0.001%. A 0.001% DMSO mock in ½ MS medium did not elicit [Ca²⁺]_cyt_ responses (n = 5 roots; kymographs and crestplots in Supplemental Data), consistent with the absence of responses to solvent controls in *UBQ10::R-GECO1* imaging (Waadt et al., 2017). Cold treatment was administered to roots as an ice-chilled ½ MS medium (approximately 0°C at application). Chitin was not prepared as a stock: chitin from shrimp shells (practical grade, Sigma-Aldrich Cat. No. C7170) was weighed directly, ground by shaking in liquid nitrogen for 30 s on an amalgamator (frequency 60 Hz), and suspended in Milli-Q H₂O at a final concentration of 100 µg mL⁻¹. Unless stated otherwise, elicitors were applied at the following final concentrations in ½ MS medium: NLP20, 10 µM; FLG22, 10 µM; C8, 10 µM; BcPG3, 5 µM; cellobiose, 100 µM; PEP1, 10 µM; eATP, 150 µM; L-Glu, 1 mM; RALF23, 10 µM; NaCl, 100 mM; KCl, 100 mM; Ca(NO₃)₂, 100 mM; KH₂PO₄, 100 mM; KNO₃, 100 mM; NH₄NO₃, 100 mM; (NH₄)₂HPO₄, 100 mM; IAA, 1 µM; D-sorbitol, 200 mM. Chitin (Fig. 1) was applied at 100 µg mL⁻¹.

The biostimulant used in this study (BC4), a proprietary Micro Carbon Technology (MCT) liquid biostimulant that contains a blend of organic molecules and colloidal minerals, was provided by Huma®, Inc. (Gilbert, AZ, USA). Its elemental composition is given in Table S1. Raw biostimulant was diluted in ½ MS medium to a final concentration of 4 µg C mL⁻¹, which raised the pH of the working solution from 5.75 to 6.03. Mock-treated roots received ½ MS medium adjusted to pH 6.03, were processed identically and without bleaching or drift correction, and showed no fingerprint.

### Root growth assay

Control plates contained ½ MS agar medium as described above. For BC4 plates, filter-sterilized BC4 was added to the ½ MS agar medium to a final concentration of 4 µg C mL⁻¹; mock plates received the same volume of sterile ½ MS medium adjusted to pH 6.03 to account for dilution of the agar medium. Seeds were sown directly on the plates and grown for 10 days under the conditions described above, after which plates were scanned on a flatbed scanner (EPSON Perfection V370) and primary root length was measured in Fiji. Experiments were performed in two independent batches (n = 14–38 roots per condition); statistical analysis is described in the legend of Fig. 7E.

### [Ca²⁺]_cyt_ imaging with high-resolution microscopy

High-resolution time-lapse imaging of *Arabidopsis* single-roots expressing the *UBQ10::R-GECO1* calcium biosensor (Fig. 1, S1) was performed using a custom-built fluorescence microscope based on a Nikon Ti-E stand. The system was equipped with a 10× objective (S Fluor, N.A.=0.50, Nikon), a motorized stage (Applied Scientific Instrumentation, USA), a motorized filter wheel (Cairn Research, UK), a laser launch system (Omicron, Germany), a triple-band dichroic mirror (Chroma 440/514/561), and an sCMOS camera (Teledyne Photometrics Prime 95B, USA). The *UBQ10::R-GECO1* biosensor line was imaged with 561 nm laser excitation at 10 mW power, using a 605/70 nm emission filter. Image acquisition was carried out using the Nikon NIS-Elements software (version 4.0), with a time resolution of 1 fps and an exposure time of 700 ms.

Post-acquisition, image data were processed and analyzed using ImageJ/Fiji (version 1.49b). The workflow included image alignment using the MultiStackRegistration plugin (https://github.com/miura/MultiStackRegistration), background subtraction (rolling ball radius = 200 px), and application of a Gaussian blur filter (sigma = 1 px).

### [Ca²⁺]_cyt_ imaging with low-resolution microscopy with the BalanceSir platform

Low-resolution imaging was performed using 7-day-old seedlings grown on a sterile nylon mesh (Schwegmann Filtration Technology, 100 µm pore size, Cat.No. SG00020140-9068289) placed on the surface of agar medium (Fig. S2A). Prior to the imaging session, the mesh with seedlings was transferred into a cell culture chamber (Sarstedt, cell culture chamber, 1-well, on a cover glass, Cat. No. 94.6190.102), with one mesh per chamber (Fig. S2C).

Chambers were plasma-treated (HAR-040 Diener/Femto Plasma Cleaner) to ensure uniform medium distribution (Fig. S2B). A mesh with 7-day-old seedlings was then placed into the well chamber, and 1 mL of ½ MS medium was added. Seedlings were equilibrated in an airtight container, in the dark, for 30 minutes prior to imaging.

Fluorescence imaging was performed using Olympus and Nikon stereo microscopes. The Olympus MVX10 stereo-microscope was equipped with a 2x objective (MVPLAPO, 0.5 NA, FOV=17 mm), a motorized focus (SZX2-FOF-M), a CoolLED pE4000 light source ( 50% 550 nm excitation), quad-band dichroic mirrors 395/25, 475/28, 555/25, 635/18 (Chroma 89402bs), a TwinCam image splitter with GFP/mApple Laser Beamsplitter H 560 LPXR superflat (AHF F48-558), a bandpass mApple emission filter (Chroma ET595/33m), and a CellCam Kikker 100MT camera in 2×2 binning mode (14bit, px=4.6744 µm, Cairn UK). Image acquisition was done through Micro-Manager software version 2.0 (Edelstein *et al*., 2014), with an exposure time of 700 ms, and images were acquired at 1fps. The Nikon SMZ18 stereo microscope was used for low-resolution imaging as well; equipped with a SHR Plan Apo 2x objective (Nikon, NA 0.3), Lumencor SOLA light engine, with a 545/25 nm excitation filter and a 605/70 nm emission filter (Chroma). Images were captured using an ORCA-Flash 4.0 sCMOS camera (Hamamatsu, Japan). Image acquisition was done through NIS Elements software (version 4.0), the exposure time was set to 700 ms, and the images were acquired at 1fps.

Before imaging began, the well chamber was mounted onto the BalanceSir platform (tilting device, Fig. S2E), and the focal plane containing multiple roots was adjusted (Fig. S2G, Supplemental Movie 2). Once imaging commenced, a baseline recording was acquired for up to 5 min, after which the ½ MS medium was replaced with medium containing the elicitor (Supplemental Movies 1 and 2). Imaging continued for 15 min following elicitor application, after which a positive control treatment with 150 µM eATP was applied to confirm biosensor functionality and responsiveness.

Ca²⁺tlas recordings were acquired on both systems, and biostimulant and mutant recordings on the Olympus MVX10. The two systems have closely matched effective pixel sizes (Nikon 4.33 µm, Olympus 4.67 µm), and wild-type FLG22 and NaCl responses recorded on the Olympus alongside the mutant experiments fell within the range of the Ca²⁺tlas data for all NaCl descriptors and for four of six FLG22 descriptors; these FLG22 signatures initiated further from the root tip and spread further, but occupied the same region of signaling space as the Ca²⁺tlas FLG22 responses (Fig. S5D,E).

### Time-series processing and kymograph generation from low-resolution imaging

Once generated, the time-series data were used to create space-time plots, hereafter referred to as kymographs. Image processing was performed using ImageJ/Fiji (version 1.49b). After an initial inspection of the image stack during baseline and positive control imaging for clarity, the treatment section was extracted and used for further processing. Each root was defined with a centerline ROI of 5-pixel width (Fig. S2G), and the time-series data were processed using the “ROI into kymographs.ijm” macro to generate “raw” kymographs.

The kymographs were further processed with the “Crestline batch.ijm” macro to enhance the signal-to-noise ratio in the raw kymographs, resulting in normalized kymographs, hereafter referred to as crestplots. Signal normalization was performed separately for each position along the root using min–max scaling over the entire recording, according to (y − min(x)) / (max(x) − min(x)), where y is the intensity at a given time point and x is the set of intensity values at that position over time; this standardizes local intensity dynamics while preserving relative spatiotemporal features. Since this processing step leads to a loss of pixel size metadata, it was necessary to re-embed the metadata into both the kymographs and crestplots using the “Change Properties Batch.ijm” macro. All macros are available as part of the supplemental data. Re-embedded pixel-size metadata were verified against the original acquisition calibration prior to descriptor extraction, ensuring that all distance and velocity measurements retained the correct spatial scaling. All analysis steps were performed using a standardized pipeline to ensure reproducibility across experiments.

### Feature extraction

Ca²⁺ fingerprint dynamics were quantified manually in Fiji/ImageJ (Schindelin et al., 2012) by measuring six descriptors on crestplots with the rectangle selection tool (Fig. 4). Time was measured from the return of the BalanceSir platform to the imaging position, which marks the start of treatment (t = 0). Signal onset time (SOT) was defined as the time of the first clearly visible signal that could be traced into a continuous fingerprint, and its position gave the spatial origin, expressed as distance from the root tip (DfRT). Signal duration (SD) was the temporal extent of the initial transient, and signal spatial spread (SSS) the length of root covered by the fingerprint from its origin. Tipward and shootward velocities (Vt and Vs) were calculated from a rectangle fitted to the respective fingerprint front, as the ratio of its spatial to its temporal extent. Because onset was identified visually rather than by a numerical threshold, the approach was validated by three independent blinded scorers who extracted descriptor values from a shared subset of crestplots, yielding highly comparable results across observers. Descriptors were extracted from full-length kymographs covering the entire imaged root segment (approximately 3–9 mm, depending on the recording); crestplots shown in the figures are cropped to the first 3.4 mm, which spans the meristematic, elongation and differentiation zones, for consistent display. Spatial spread was measured within the imaged segment and is therefore a lower bound where a fingerprint extended beyond it. Crestplots with imaging artifacts (e.g., drift, focus loss or root movement) or with a discontinuous signal front that could not be traced unambiguously along the root were excluded from parameterization. All kymographs and crestplots, including excluded recordings, are deposited at Zenodo.

### Data analysis

Quantitative data were analyzed, tested, and plotted with R version 4.3.2 and R Studio version 2024.12.1 build 563 (R Core Team, 2020). Each elicitor, genotype and treatment combination was recorded in at least three independent imaging sessions, and individual roots were treated as biological replicates. In the reporter-line experiments (Fig. S8), roots without a fingerprint that could be parameterized were counted as non-responders and excluded from descriptor statistics; response rates (responding/total roots) were compared by Fisher’s exact test. Data were analyzed by comparing differences between treatments using a Kruskal-Wallis test, followed by Dunn’s post hoc test with Benjamini-Hochberg correction (FSA package). Significant differences were visualized using compact letter displays (multcompView package), based on adjusted p-values. Compact letter displays were arranged such that ‘a’ corresponds to the highest median value. Boxplots were generated using the “ggplot2” package (http://ggplot2.tidyverse.org/).

### PCA projection and posterior class membership of supplementary individuals

PCA was performed on the six untransformed quantitative Ca²⁺ response descriptors (SOT, DfRT, SSS, SD, Vt, and Vs) after centering and scaling using FactoMineR (v2.10). Velocities were used untransformed; all 561 reference roots had non-zero tipward and shootward velocities, so no values were missing. Biplots were produced with a factoextra package (Kassambara and Mundt, 2020). Reference elicitor recordings (Ca^2+^tlas Panel) were treated as active individuals and used to estimate PCA loadings and axes. Biostimulant parameters were treated as supplementary individuals and were projected *post-hoc* into the fixed PCA space (i.e., they did not contribute to estimation of PCA axes or class ellipses). Class labels for active samples were defined a priori: MAMP (NLP20, FLG22, C8, BcPG3), DAMP (cellobiose, eATP, L-Glu), signaling (PEP1, RALF23, IAA), nutrients/salts (NaCl, KCl, Ca(NO₃)₂, KH₂PO₄, KNO₃, NH₄NO₃, (NH₄)₂HPO₄), osmotic stress (D-sorbitol) and temperature stress (cold).

Posterior membership of supplementary individuals to the pre-defined elicitor classes was estimated in PCA space under class-conditional MultiVariate Normal (MVN) models. For each class, the mean vector and covariance matrix were estimated from active samples in PCA space, with covariance shrinkage performed using corpcor (v1.6.10; cov.shrink) to improve numerical stability. Class likelihoods were computed using mvtnorm (v1.2-4), and posterior probabilities were obtained via Bayes’ rule using class priors proportional to class frequencies among active samples. Because class frequencies in the reference panel reflect sampling effort rather than the probability that an unknown stimulus belongs to a given class, BC4 posteriors were additionally computed with uniform class priors (Table S3). BC4 quantitative calcium responses of *moca1 UBQ10::R-GECO1 #2-1* and *cerk1-2 UBQ10::R-GECO1 #1-3* were projected into the same PCA space as supplementary individuals and evaluated with the same class models; genotypes were compared by Kruskal–Wallis tests followed by Dunn’s post hoc test with Benjamini–Hochberg correction (Fig. S11; Table S5). Unless stated otherwise, inference was performed using the first K principal components capturing ≥80% of the total variance (K = 4 in this study; 87.4% variance explained). Posterior probabilities were visualized as stacked barplots (per-individual membership) and cumulative posterior mass curves (class-level support).

### Posterior model validation

To quantify class assignment within the signaling space, we implemented a probabilistic classification framework. Model performance was evaluated on the active reference elicitors using Stratified K-Fold Cross-Validation (SKFCV, n=5). In each fold, samples were partitioned to approximately preserve class proportions, the class-conditional MVN parameters and class priors were re-fit using the training split only, and posterior probabilities were computed for the held-out split in the same PCA space (first K = 4 PCs). Predicted labels were assigned by maximum posterior probability. Performance was summarized by the confusion matrix (row-normalized), overall accuracy, and balanced accuracy (mean per-class recall, i.e., the proportion of samples from a given class correctly assigned to that class), with class-wise recall values reported in Table S2. To test prediction for stimuli, absent from the reference panel, we additionally performed leave-one-elicitor-out validation, in which all replicates of one elicitor were withheld, the PCA and class models were refit on the remaining elicitors, and the withheld elicitor was projected and assigned to the class predicted for most of its replicates, under both frequency-based and uniform class priors. Classes represented by a single elicitor (osmotic and temperature stress) could not be evaluated in this way (Table S4).

### Correlation analysis

To characterize how Ca²⁺ fingerprint descriptors co-vary within each stimulus condition, we computed within-treatment Pearson correlations across the six quantitative response variables (SOT, DfRT, SSS, SD, Vt, and Vs). For each treatment, correlations were calculated using complete-case observations (rows with missing values removed) using Hmisc (rcorr) and extracted as correlation coefficients (r). P-values were computed by rcorr and adjusted within treatment using the Benjamini–Hochberg procedure; however, correlation maps were used primarily as descriptive summaries of parameter coupling rather than for significance thresholding. To visualize the treatment-specific correlation structure compactly, correlation coefficients for each descriptor pair (upper triangle only) were displayed as a mosaic heatmap in which each treatment was represented as a sub-tile offset within the corresponding variable–variable cell (tile fill = Pearson’s *r*), implemented in ggplot2.

## Data availability

The Fiji macros of the Crestline pipeline are archived at Zenodo (DOI: 10.5281/zenodo.23182873). The R script that reproduces all statistical and multivariate analyses from Supplemental Datasets 1–10 is available at https://github.com/milanzupunski/PlantCalciumAtlas. The quantitative descriptor data underlying the Ca²⁺tlas and the biostimulant experiments are deposited at Zenodo (DOI: 10.5281/zenodo.23183343), and the corresponding kymographs and crestplots at Zenodo (DOI: 10.5281/zenodo.23184086). Any additional data are available from the corresponding authors upon request.

## Author contributions

Conceptualization: G.G., J.P.

Methodology: M.Ž., R.B., M.Z., G.G., J.P., K.L. (BalanceSir platform), R.W., M.K. (Ca²⁺ imaging and image processing)

Investigation: M.Ž., R.B., M.Z., M.G., R.L., K.L.

Resources: R.W., K.S. (R-GECO1 plasmids and plants); R.L. (biostimulant product)

Formal analysis: M.Ž., R.B., M.Z., M.G.

Writing – original draft: M.Ž., G.G.

Writing – review & editing: all authors

Supervision: G.G., J.P.

Funding acquisition: G.G., J.P., M.Ž.

## Acknowledgements

The authors thank all former and current members of the Grossmann Lab for insightful discussions, Michaela Gerads and Jacqueline Esser for technical assistance, and Claudia Franken-Stemmler for administrative support. We are also grateful to Prof. Thorsten Nürnberger (University of Tübingen, Germany) for providing bacterial peptide elicitors; to Prof. Dr. Zhonghao Jiang (Shenzhen University, China) for providing *moca1* seeds.

## Declaration of Generative AI and AI-assisted technologies in the writing process

During the preparation of this work the authors used Claude (Anthropic) in order to consolidate and document the R analysis code, and improve the language and readability of the text. After using this tool, the authors reviewed and edited the content as needed and take full responsibility for the content of the publication.

## Supporting Information

**Figure S1 *UBQ10::R-GECO*1 Ca²⁺ signal amplitude and apparent spatiotemporal dynamics depend on ROI placement.**

All panels derive from the same two recordings shown in Fig. 1, one root treated with 150 µM eATP (left column) and one with 100 µg mL⁻¹ chitin (right column), analyzed with a centerline ROI (a–b) or an edge-positioned ROI (c–d), both 5 pixels wide. A-B) Centerline ROI. (a) Root-tip images showing the ROI position used for intensity extraction (scale bar, 50 µm) and the resulting ΔF/F₀ traces. (b) Corresponding kymographs (royal LUT) displaying Ca²⁺ signal intensity as a function of time (x-axis) and position along the root (y-axis, root tip at the bottom; 1 pixel column = 1 s). Corresponding crestplots (wave_tracking LUT) derived from the same recordings, highlighting the extracted fingerprint trajectories. C-D) Edge-positioned ROI. (c) ROI placement for intensity extraction (scale bar, 50 µm) and the resulting ΔF/F₀ traces. (d) Kymographs (royal LUT); Crestplots (wave_tracking LUT). Axes in (D) are as in (B). The royal LUT spans a fixed display range (min 100 to max 800, arbitrary units) applied identically to all panels, so intensities are directly comparable between them. The wave_tracking LUT spans the full 16-bit output range of the Crestline normalization (0 to 65,535) in all panels; values are normalized over time at each root position, so they represent relative signal within each position’s time course rather than absolute [Ca²⁺]cyt. Together, the comparisons show that ROI placement (centerline vs. edge) influences both the measured fluorescence amplitude and the inferred spatiotemporal fingerprint motivating standardized ROI definitions for quantitative analyses.

**Figure S2 BalanceSir imaging platform and seedling growth on nylon mesh.**

A) Seedlings grown for 7 days on nylon mesh placed on top of ½ MS agar. B) Plasma treatment of imaging chambers (HAR-040 Diener/Femto Plasma Cleaner, Germany) to reduce surface tension and improve wetting during imaging. C) Transfer of the mesh with seedlings into the imaging chamber followed by addition of 1 mL ½ MS liquid medium. D) Top view of the chamber containing the mesh. The orthogonal fiber layout guides roots to grow straight and in parallel, enabling simultaneous imaging of multiple roots within a single field of view. E) Chamber mounted on the BalanceSir platform for imaging. F) Tilting of the BalanceSir platform to facilitate rapid exchange of the liquid medium by pipetting without disturbing the root positioning. G) Imaging field of view (FOV) of roots grown on nylon mesh. The magnified inset illustrates typical ROI placement used for downstream image analysis.

**Figure S3 Representative kymographs of stimulus-evoked Ca²⁺ dynamics in Arabidopsis roots expressing UBQ10::R-GECO1 for each tested elicitor.**

Each elicitor and its concentration are indicated above the corresponding panel. The x-axis denotes time (1 pixel column = 1 s), and the y-axis denotes position along the imaged root segment (root tip at the bottom). For elicitors with known perception pathways, the corresponding receptor or sensing component expression profile is shown next to each kymograph (data from the BAR ePlant resource), with the relevant signaling receptor indicated in parentheses; for elicitors with unknown receptors, a grayscale root image is shown instead. The royal LUT spans a fixed display range (min 100 to max 800, arbitrary units) applied identically in all panels, so intensities are directly comparable between them. Expression profiles are shown as relative expression values on the adjacent color scale. They are published reference data (Waese et al., 2017), scaled to match the developmental zonation of the imaged roots using the brightfield image in Fig. 2 as reference; they were not measured in the imaged roots and are shown for exploratory comparison only. Corresponding crestplots are shown in Fig. 2.

**Figure S4 Quantitative spatiotemporal descriptors distinguish stimulus-specific Ca²⁺ fingerprints in Arabidopsis roots.**

A) Delay from stimulus application to first detectable Ca²⁺ signal (onset time). B) Distance from the root tip at which the first signal was detected. C) Duration of the initial Ca²⁺ signal. D) Spatial spread of the signal from its point of origin. E) Tipward propagation velocity (Vt; toward the root tip). F) Shootward propagation velocity (Vs; away from the root tip). In (e) and (f), velocities are plotted on a logarithmic axis for display only; untransformed values were used in all statistical and multivariate analyses. Elicitors are ordered by stimulus class: MAMPs (NLP20, FLG22, C8, PG3), DAMPs (D-Cellobiose, PEP1, eATP, L-Glu), nutrients/salts (NaCl, KCl, Ca(NO₃)₂, KH₂PO₄, KNO₃, NH₄NO₃, (NH₄)₂HPO₄), signaling ligands (RALF23, IAA; PEP1 was assigned to this class in the multivariate analyses), osmotic stress (D-Sorbitol), and temperature stress (Cold 1/2MS). Data are shown as boxplots (median + IQR with whiskers (1.5×IQR)) with individual replicates overlaid. Omnibus differences among treatments were significant for all six descriptors (Kruskal–Wallis, df = 18: onset time H = 481.0, p = 1.0 × 10⁻⁹⁰; distance from tip H = 349.9, p = 2.4 × 10⁻⁶³; duration H = 384.0, p = 1.9 × 10⁻⁷⁰; spatial spread H = 333.8, p = 5.1 × 10⁻⁶⁰; tipward velocity H = 461.5, p = 1.2 × 10⁻⁸⁶; shootward velocity H = 466.2, p = 1.3 × 10⁻⁸⁷). Different letters denote significant differences among treatments (Dunn’s post hoc test with Benjamini–Hochberg correction, p < 0.05). Replicate numbers: NLP20 = 14; FLG22 = 79; C8 = 17; PG3 = 44; cellobiose = 17; PEP1 = 30; eATP = 29; L-Glu = 28; RALF23 = 20; NaCl = 61; KCl = 25; Ca(NO₃)₂ = 27; KH₂PO₄ = 15; KNO₃ = 22; NH₄NO₃ = 23; (NH₄)₂HPO₄ = 32; IAA = 24; D-sorbitol = 21; cold (½ MS) = 33.

**Figure S5 Robustness of Crestline fingerprints across biosensors and imaging platforms.**

A) Representative kymograph (left) and corresponding crestplot (right) of cytosolic Ca²⁺ dynamics in *Arabidopsis* roots expressing *35S::GCaMP3* following elicitation with 150 µM eATP (DORN1/P2K1-dependent; n = 8). B) As in (A), following elicitation with 1 mM L-glutamate (GLR-dependent; n = 8). In (A) and (B), the x-axis denotes time (1 pixel column = 1 s), and the y-axis denotes position along the root (root tip at the bottom). The royal LUT spans a fixed display range (min 100 to max 800, arbitrary units) applied identically to both kymographs, so intensities are directly comparable between them. The wave_tracking LUT spans the full 16-bit output range of the Crestline normalization (0 to 65,535), applied identically to both crestplots; values are normalized over time at each root position, so they represent relative signal within each position’s time course rather than absolute [Ca²⁺]_cyt_. C) PCA of the six Ca²⁺ fingerprint descriptors fitted on *UBQ10::R-GECO1* recordings for 150 µM eATP (n = 29) and 1 mM L-Glu (n = 28) from the Ca²⁺tlas. *35S::GCaMP3* replicates (eATP n = 9, L-Glu n = 8; recorded on the Olympus MVX10) are projected as supplementary individuals and do not contribute to the PCA. Open symbols mark R-GECO1 group centroids; ellipses show 95 % confidence ellipses of the R-GECO1 groups. D) Fingerprint descriptors of wild-type *UBQ10::R-GECO1* roots treated with 10 µM FLG22 or 100 mM NaCl, recorded for the Ca²⁺tlas (Nikon SMZ18; n = 79 and 61) or alongside the mutant experiments (Olympus MVX10; n = 23 and 11); Mann–Whitney tests, Benjamini–Hochberg-adjusted; the larger field of view of the Olympus allows signals to be followed over a longer root segment; velocities on a log axis for display only. E) PCA fitted on the Ca²⁺tlas FLG22 and NaCl roots (filled symbols, solid 95 % normal ellipses), with Olympus roots projected as supplementary individuals (crosses, dashed ellipses).

**Figure S6 How closely individual biostimulant replicates match each reference stimulus category, and how accurate this classification approach is.**

A)Representative kymographs (royal LUT) of cytosolic Ca²⁺ dynamics in wild-type Arabidopsis root tips expressing R-GECO1 following mock/biostimulant treatment. the x-axis denotes time (1 pixel column = 1 s), and the y-axis denotes position along the root (root tip at the bottom). The royal LUT spans a fixed display range (min 100 to 800, arbitrary units) applied identically to both kymographs, so intensities are directly comparable between them. B) Per-individual posterior membership of biostimulant supplementary individuals to the pre-defined elicitor classes, displayed as a stacked barplot (individuals ordered by maximum posterior probability). Under frequency-based priors, posterior support is split mainly between the Nutrients/Salts and DAMP classes, whereas MAMP and osmotic/temperature-stress classes contribute little. C) The same posterior probability matrix shown as a heatmap (rows = biostimulant data; columns = elicitor classes; colour indicates posterior probability), highlighting the consistency of class support across individuals and potential mixed-class assignments. D) Stratified 5-fold cross-validation of the posterior class-conditional multivariate normal (MVN) classifier on the active reference elicitor dataset. The confusion matrix reports row-normalized fractions (colour scale) with raw counts overlaid in each cell. Overall accuracy (acc = 0.74) and balanced accuracy (bal.acc = 0.68) are indicated in the title. Consistent with the diagonal structure, MAMP and nutrients/salts show high recall, while DAMP exhibits lower recall (greater overlap with other classes); corresponding class-wise recall values are provided in Table S2. E) Quantitative spatiotemporal descriptors of Ca²⁺ fingerprints in *Arabidopsis* roots treated with the biostimulant. Boxplots show signal onset time, distance from the root tip, signal duration, spatial spread, and tipward/shootward velocities for individual roots. Dashed lines indicate the median response of representative Nutrients/Salts (NaCl) and DAMPs (eATP) treatments for comparison.

**Figure S7 Functional validation of R-GECO1 Ca²⁺ reporter lines in mutant backgrounds.**

Spatiotemporal Ca²⁺ dynamics in *Arabidopsis* roots expressing *UBQ10::R-GECO1* in wild type (Col-0) and in two independent reporter lines for each mutant background (*moca1*, *cerk1-2*) were recorded following elicitation with 100 mM NaCl (a,b) or 10 µM FLG22 (c,d), followed by 150 µM eATP as a positive control. For each genotype, kymographs and corresponding crestplots are shown (a,c), together with ΔF/F₀ traces (mean ± SD across replicates) aligned to stimulus application (vertical dashed lines) (b,d). F₀ is the mean of the 60 s pre-stimulus baseline; for the eATP phase, F₀ is the mean of the last 20 s before eATP application. (a,b) NaCl responses were used to verify the expected salt-induced Ca²⁺ fingerprint phenotype in each background. The black dashed line in (b) indicates the mock treatment (½ MS medium). (c,d) FLG22, reported to act independently of the tested mutant backgrounds, was used as a second validation stimulus; all lines exhibited a clear FLG22-induced Ca²⁺ response. In kymographs and crestplots, the x-axis denotes time (1 pixel column = 1 s), and the y-axis denotes position along the root (root tip at the bottom left). The royal LUT spans a fixed display range (min 100 to max 800, arbitrary units) applied identically to all kymographs, so intensities are directly comparable between them. The wave_tracking LUT spans the full 16-bit output range of the Crestline normalization (0 to 65,535), applied identically to all crestplots; values are normalized over time at each root position, so they represent relative signal within each position’s time course rather than absolute [Ca²⁺]_cyt_.

**Figure S8 Fingerprint descriptors of FLG22 and NaCl responses in the reporter lines.**

(a) NaCl (100 mM) and (b) FLG22 (10 µM) responses of wild type (Col-0) and two independent UBQ10::R-GECO1 reporter lines each of moca1 (#1-1, #2-1) and cerk1-2 (#1-2, #1-3), from the experiments shown in Fig. S7. Numbers in parentheses give responding/total roots; non-responding roots are not plotted and were excluded from descriptor statistics. Kruskal–Wallis test followed by Dunn’s post hoc test with Benjamini–Hochberg correction on responding roots; different letters denote significant differences (p < 0.05). NaCl response rates differed among lines (Fisher’s exact test, p < 0.0001). Velocities are shown as log₁₀ for display only; untransformed values were used for statistics.

**Figure S9 Kymographs, intensity traces and fingerprint descriptors of BC4-evoked Ca²⁺ signatures in wild type, *moca1* and *cerk1-2*.**

A) Representative crestplots (top) and kymographs (bottom) of cytosolic Ca²⁺ dynamics in *Arabidopsis* roots expressing *UBQ10::R-GECO1* in wild type (Col-0), *cerk1-2* (#1-3) and *moca1* (#2-1) backgrounds (left to right) following mock treatment (½ MS medium). B) Representative kymographs for the same genotypes following biostimulant treatment (BC4, 4 µg C mL⁻¹ in ½ MS medium). In (a) and (b), the x-axis denotes time (1 pixel column = 1 s), and the y-axis denotes position along the root (root tip at the bottom). The royal LUT spans a fixed display range (min 100 to max 800, arbitrary units) applied identically to all kymographs, so intensities are directly comparable between them. The wave_tracking LUT spans the full 16-bit output range of the Crestline normalization (0 to 65,535), applied identically to all crestplots; values are normalized over time at each root position, so they represent relative signal within each position’s time course. C) Corresponding ΔF/F₀ traces (mean ± SD) aligned to stimulus application (vertical dashed lines) for the sequential treatment ½ MS → mock or biostimulant → 150 µM eATP, providing the amplitude-based readout underlying the crestplots in Fig. 7A. The dashed black line indicates mock, and the magenta line indicates biostimulant treatment. F₀ is the mean of the 60 s pre-stimulus baseline; for the eATP phase, F₀ is the mean of the last 20 s before eATP application. D-G) Fingerprint descriptors not shown in Fig. 7: D) signal onset time, E) distance from the root tip at first detection, F) tipward and G) shootward velocity (log axis for display only). n = 20 (Col-0), 13 (*moca1* #2-1) and 12 (*cerk1-2* #1-3) roots. Letters denote groupings from Dunn’s post hoc test with Benjamini–Hochberg correction (p < 0.05).

**Figure S10 Primary root growth under BC4 treatment.**

A) Representative primary root growth phenotypes on ½ MS agar under control (½ MS), mock and BC4 (4 µg C mL⁻¹) treatment for wild type, cerk1-2 and moca1 (left to right). Scale bar = 10 mm. B) BC4 treatment did not significantly affect primary root length in the *cerk1-2* (#1-3) and *moca1* (#2-1) *UBQ10::R-GECO1* reporter lines, consistent with Fig. 7E. Two independent batches; n = 17–33 roots per condition.

**Figure S11 BC4 responses of *moca1* and *cerk1-2* projected into the Ca²⁺tlas signaling space.**

A) Reference stimulus classes (95% data ellipses, as in Fig. 6b) with BC4-treated roots of wild type (magenta asterisks), *moca1 UBQ10::R-GECO1* #2-1 (teal triangles) and *cerk1-2 UBQ10::R-GECO1* #1-3 (gold squares) projected as supplementary individuals; diamonds mark genotype centroids. B) Squared Mahalanobis distance of each BC4-treated root to the nutrient/salt class centroid in the first four principal components. C) Posterior probability of nutrient/salt class membership under frequency-based priors. In B) and C), Kruskal–Wallis test followed by Dunn’s post hoc test with Benjamini–Hochberg correction; different letters denote significant differences (p < 0.05). n = 20 (wild type), 13 (*moca1*) and 12 (*cerk1-2*) roots. Data in Supplemental Datasets 3 and 4; statistics in Table S5.

**Supplemental Movie 1 Medium exchange on the BalanceSir tilting platform during live root Ca²⁺ imaging.**

Real-time demonstration of the imaging workflow on the microscope. Seedlings are imaged under ½ MS medium to establish a baseline, the BalanceSir platform is manually tilted to pool liquid away from the root tips. ½ MS medium is removed by pipetting and replaced by the treatment medium, followed by the platform being returned to the imaging position to resume acquisition without disturbing the root placement.

**Supplemental Movie 2 Live acquisition view during tilting and eATP-triggered Ca²⁺ signaling.**

Screen capture of the camera output corresponding to the medium-exchange workflow in Movie 1. The video is shown in false color (mpl_inferno LUT) acquired at 1 frame per second and played back at 30 frames per second. Baseline fluorescence is followed by a transient focal-plane shift during platform tilting and restoration of the original focus after tilt-back. The timestamp is set to 00:00 at the moment the platform returns to the imaging geometry, which marks the start of the 150 µM eATP treatment. Within seconds, root tips respond with a rapid local Ca²⁺ increase in the root tip, followed by a pronounced travelling Ca²⁺ signature approximately 1 min after stimulation. Scale bar = 1 cm.

**Table S1 Elemental profile of the biostimulant (BC4) used in this study.**

Elemental and physicochemical composition of the biostimulant stock (lot 6-23-25) as reported by fertilizer-grade analytical testing. Macro-nutrients are reported as % (w/w), trace metals as ppm, pH is measured according to EPA 150.1, and specific gravity according to ASTM D891-09. ND, not detected (below the method detection limit). Note (experimental working concentration): For imaging assays, the biostimulant was diluted to 4 µg C mL⁻¹ (4 mg L⁻¹ total C) – see Material and Methods section for the protocol. Based on the measured total carbon content (0.060% w/w) and specific gravity (1.016), this corresponds to an approximate 1:152 dilution of the stock. At this dilution, the formulation delivers approximately 1.3 mM total P (from 1.38% w/w P₂O₅, equivalent to ≈0.65 mM P₂O₅), ∼2.4 mM total N (from 0.51% w/w N, not necessarily all ionic), ∼0.24 mM K⁺ (from 0.17% w/w K₂O), ∼0.08 mM Na⁺ (from 0.028% w/w Na) and ∼3 µM Al (from 12.5 ppm), in addition to the ½ MS background medium.

**Table S2 Posterior MVN classifier performance by elicitor class (stratified 5-fold cross-validation).** Classification performance of the class-conditional multivariate normal (MVN) posterior model evaluated on the active reference elicitor dataset. For each elicitor class, the table reports the number of true samples (n) and the true-positive rate (recall), defined as the fraction of samples from that class correctly assigned to the same class (maximum posterior).

**Table S3 BC4 class assignment under frequency-based and uniform class priors.** For each reference class: prior probability under frequency-based priors, mean posterior probability across the 20 BC4 replicates under each prior, and number of replicates with maximum posterior in that class. *Legend: Frequency-based priors are proportional to class size in the reference panel; uniform priors are 1/6 for each class. Mean posteriors are averaged over the 20 BC4 replicates; replicates are assigned to the class with maximum posterior probability. Nearest: number of BC4 replicates with minimum Mahalanobis distance to that class. Median percentile: for each replicate, the fraction of the class’s own reference roots closer to the class centroid than the replicate; median over the 20 replicates (low values indicate a typical member). Within central 95%: number of replicates with a percentile below 0.95. Mean kNN share: fraction of each replicate’s 23 nearest reference roots (Euclidean distance, first four PCs) belonging to the class, averaged over replicates*.

**Table S4 Leave-one-elicitor-out validation.** For each held-out elicitor: true class, number of roots, and, under frequency-based and uniform class priors, the most frequently predicted class, the fraction of roots assigned to the true class, and the mean posterior probability of the true class. D-sorbitol and cold could not be evaluated because each is the only member of its class.

*Legend: All replicates of one elicitor were withheld, the PCA and class models were refit on the remaining elicitors with frequency-based or uniform priors, and the withheld elicitor was projected and assigned to the class predicted for most of its replicates. Twelve of 17 evaluable elicitors were assigned to their correct class under frequency-based priors (root-level accuracy 0.58), and 11 of 17 under uniform priors (root-level accuracy 0.56). D-sorbitol and cold could not be evaluated because each is the only member of its class*

**Table S5 Position of BC4 responses in the Ca²⁺tlas signaling space by genotype.**

BC4-treated roots of wild type (n = 20), *moca1* #2-1 (n = 13) and *cerk1-2* #1-3 (n = 12) projected into the reference PCA (first four PCs). (a) Mean posterior class membership (frequency-based / uniform priors). (b) Median squared Mahalanobis distance to each class centroid.

