## Supplementary material for "A calcium-fingerprint atlas infers the signaling modes of complex elicitors and reveals a salt-like response to a humic biostimulant": (Fig. S1A,D)

**This PDF file includes:**

Supplemental Figures S1–S11

Supplemental Tables S1–S5

Other supplemental information for this manuscript:

Supplemental Datasets 1–10 (Supplemental_Datasets.xlsx; Zenodo, DOI: 10.5281/zenodo.23183343)

Kymographs and crestplots (Zenodo, DOI: 10.5281/zenodo.23184086)

Fiji macros of the Crestline pipeline (Zenodo, DOI: 10.5281/zenodo.23182873)

R analysis script (https://github.com/milanzupunski/PlantCalciumAtlas)

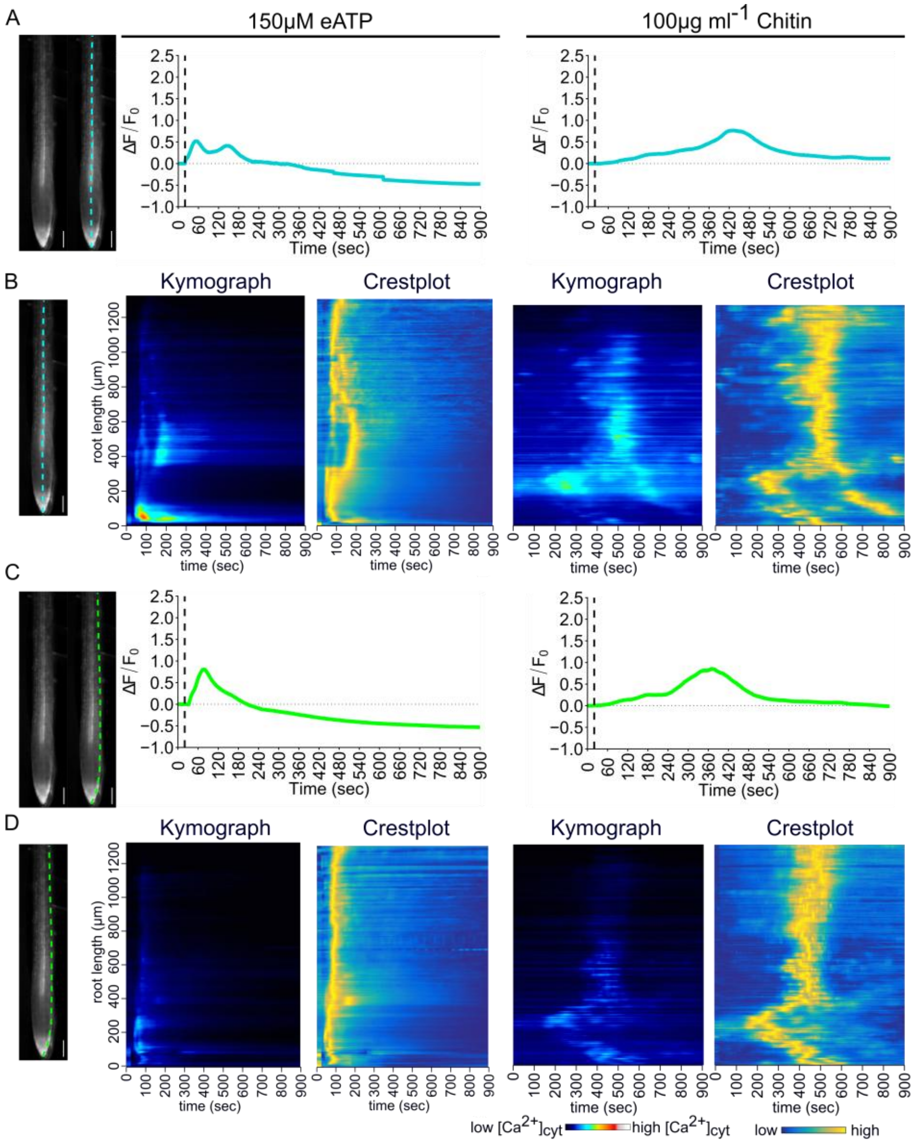


**Figure S1** ***UBQ10::R-GECO*1 $Ca^{2+}$ signal amplitude and apparent spatiotemporal dynamics depend on ROI placement.** All panels derive from the same two recordings shown in Fig. 1, one root treated with 150 µM eATP (left column) and one with 100 µg $mL^{-1}$ chitin (right column), analyzed with a centerline ROI (a–b) or an edge-positioned ROI (c–d), both 5 pixels wide. A-B) Centerline ROI. (a) Root-tip images showing the ROI position used for intensity extraction (scale bar, 50 µm) and the resulting $\Delta F/F_0$ traces. (b) Corresponding kymographs (royal LUT) displaying $Ca^{2+}$ signal intensity as a function of time (x-axis) and position along the root (y-axis, root tip at the bottom; 1 pixel column = 1 s). Corresponding crestplots (wave_tracking LUT) derived from the same recordings, highlighting the extracted fingerprint trajectories. C-D) Edge-positioned ROI. (c) ROI placement for intensity extraction (scale bar, 50 µm) and the resulting $\Delta F/F_0$ traces. (d) Kymographs (royal LUT); Crestplots (wave_tracking LUT). Axes in (D) are as in (B). The royal LUT spans a fixed display range (min 100 to max 800, arbitrary units) applied identically to all panels, so intensities are directly comparable between them. The wave_tracking LUT spans the full 16-bit output range of the Crestline normalization (0 to 65,535) in all panels; values are normalized over time at each root position, so they represent relative signal within each position's time course rather than absolute $[Ca^{2+}]_{cyt}$. Together, the comparisons show that ROI placement (centerline vs. edge) influences both the measured fluorescence amplitude and the inferred spatiotemporal fingerprint motivating standardized ROI definitions for quantitative analyses.

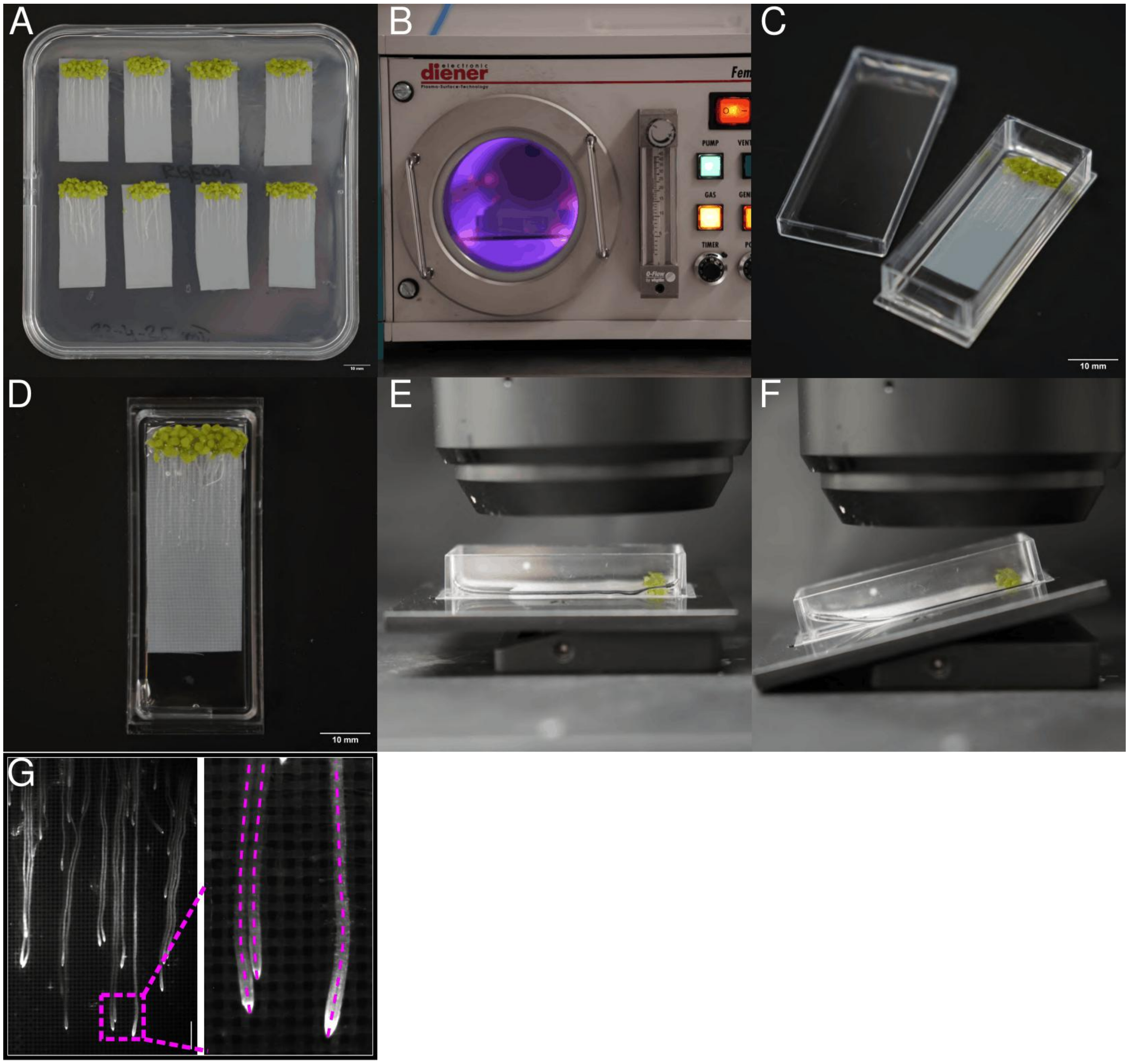


**Figure S2 BalanceSir imaging platform and seedling growth on nylon mesh.** A) Seedlings grown for 7 days on nylon mesh placed on top of ½ MS agar. B) Plasma treatment of imaging chambers (HAR-040 Diener/Femto Plasma Cleaner, Germany) to reduce surface tension and improve wetting during imaging. C) Transfer of the mesh with seedlings into the imaging chamber followed by addition of 1 mL ½ MS liquid medium. D) Top view of the chamber containing the mesh. The orthogonal fiber layout guides roots to grow straight and in parallel, enabling simultaneous imaging of multiple roots within a single field of view. E) Chamber mounted on the BalanceSir platform for imaging. F) Tilting of the BalanceSir platform to facilitate rapid exchange of the liquid medium by pipetting without disturbing the root positioning. G) Imaging field of view (FOV) of roots grown on nylon mesh. The magnified inset illustrates typical ROI placement used for downstream image analysis.

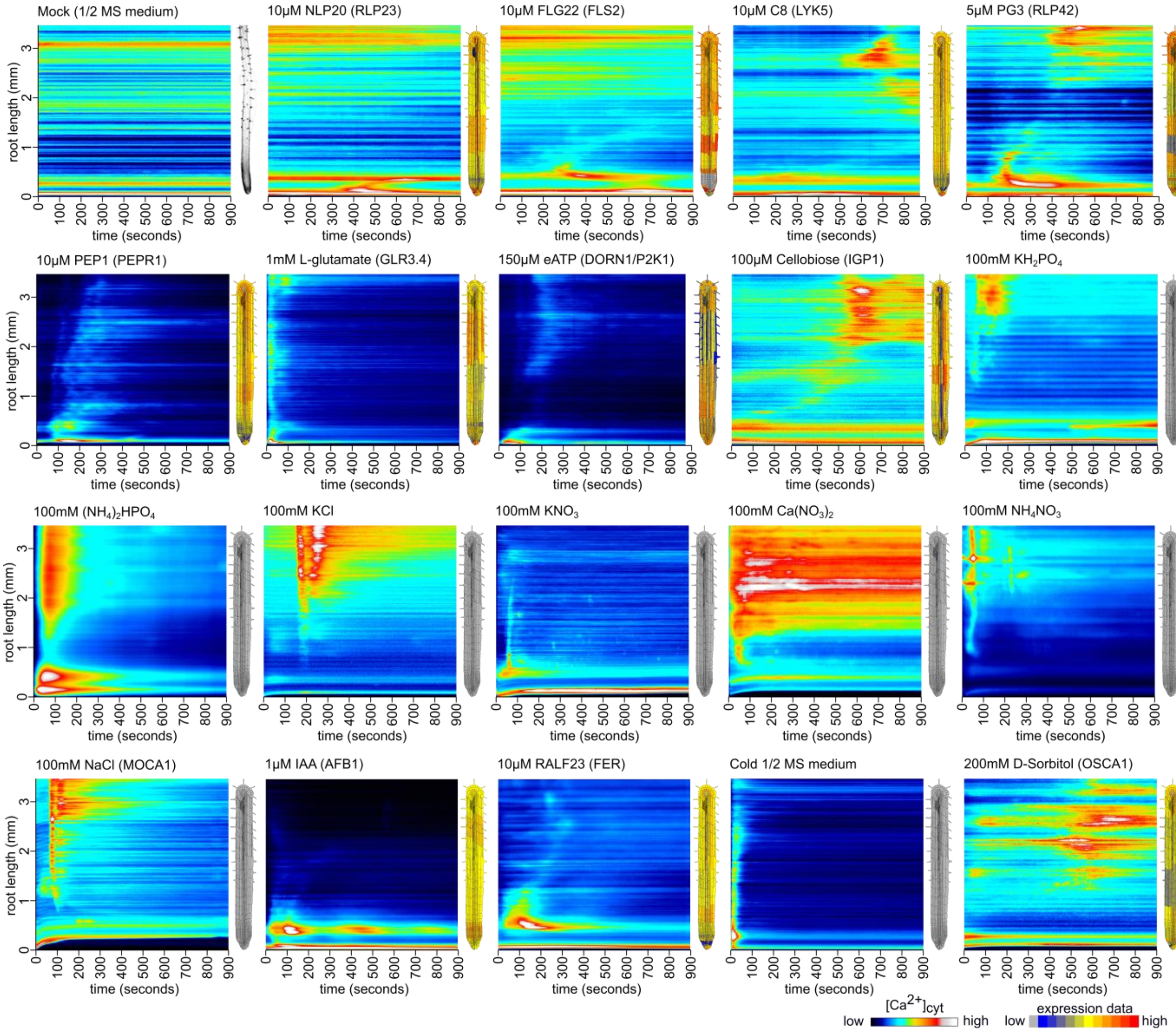


**Figure S3 Representative kymographs of stimulus-evoked $Ca^{2+}$ dynamics in *Arabidopsis* roots expressing *UBQ10::R-GECO1* for each tested elicitor.**

Each elicitor and its concentration are indicated above the corresponding panel. The x-axis denotes time (1 pixel column = 1 s), and the y-axis denotes position along the imaged root segment (root tip at the bottom). For elicitors with known perception pathways, the corresponding receptor or sensing component expression profile is shown next to each kymograph (data from the BAR ePlant resource), with the relevant signaling receptor indicated in parentheses; for elicitors with unknown receptors, a grayscale root image is shown instead. The royal LUT spans a fixed display range (min 100 to max 800, arbitrary units) applied identically in all panels, so intensities are directly comparable between them. Expression profiles are shown as relative expression values on the adjacent color scale. They are published reference data (Waese et al., 2017), scaled to match the developmental zonation of the imaged roots using the brightfield image in Fig. 2 as reference; they were not measured in the imaged roots and are shown for exploratory comparison only. Corresponding crestplots are shown in Fig. 2.

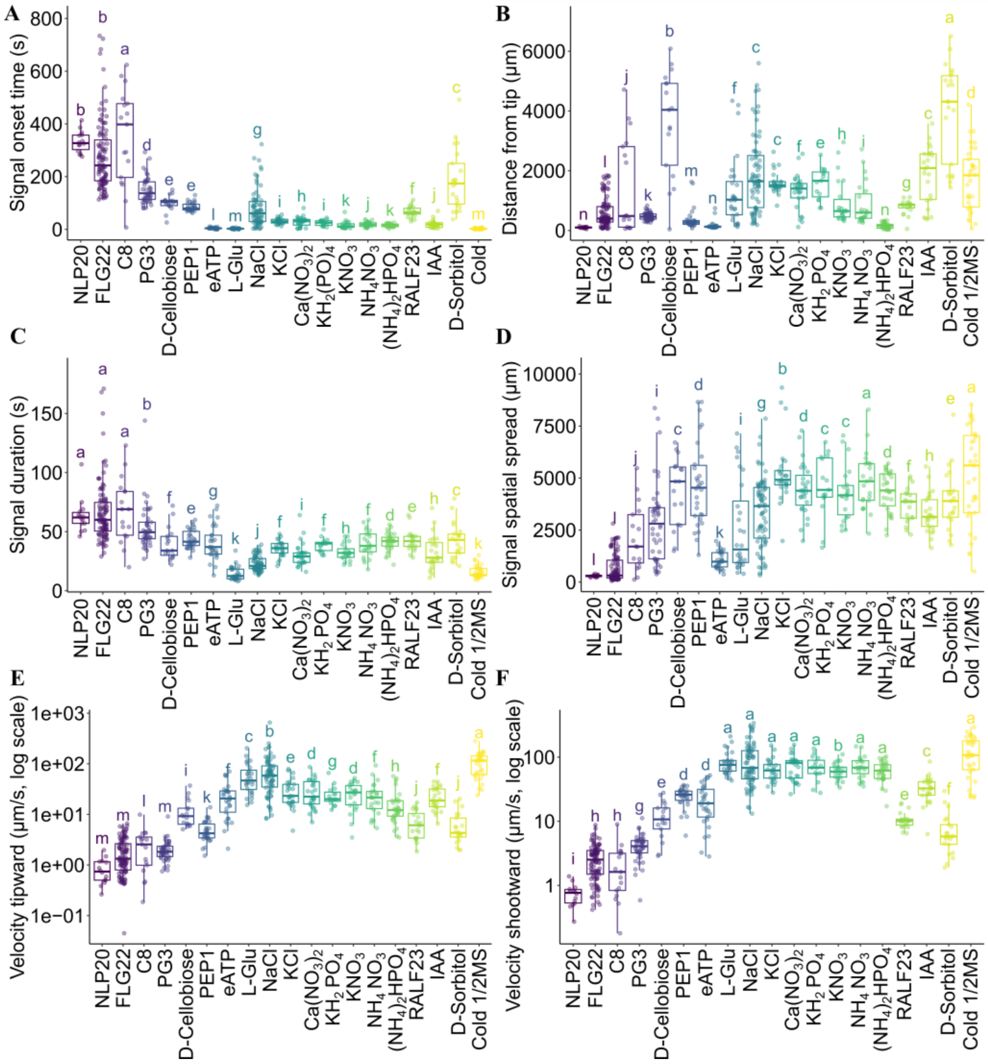


**Figure S4 Quantitative spatiotemporal descriptors distinguish stimulus-specific $Ca^{2+}$ fingerprints in *Arabidopsis* roots.** A) Delay from stimulus application to first detectable $Ca^{2+}$ signal (onset time). B) Distance from the root tip at which the first signal was detected. C) Duration of the initial $Ca^{2+}$ signal. D) Spatial spread of the signal from its point of origin. E) Tipward propagation velocity (Vt; toward the root tip). F) Shootward propagation velocity (Vs; away from the root tip). In (e) and (f), velocities are plotted on a logarithmic axis for display only; untransformed values were used in all statistical and multivariate analyses. Elicitors are ordered by stimulus class: MAMPs (NLP20, FLG22, C8, PG3), DAMPs (D-Cellobiose, PEP1, eATP, L-Glu), nutrients/salts (NaCl, KCl, $Ca(NO_3)_2$, $KH_2PO_4$, $KNO_3$, $NH_4NO_3$, $(NH_4)_2HPO_4$), signaling ligands (RALF23, IAA; PEP1 was assigned to this class in the multivariate analyses), osmotic stress (D-Sorbitol), and temperature stress (Cold 1/2MS). Data are shown as boxplots (median + IQR with whiskers (1.5×IQR)) with individual replicates overlaid. Omnibus differences among treatments were significant for all six descriptors (Kruskal–Wallis, df = 18: onset time $H = 481.0$, $p = 1.0 \times 10^{-90}$; distance from tip $H = 349.9$, $p = 2.4 \times 10^{-63}$; duration $H = 384.0$, $p = 1.9 \times 10^{-70}$; spatial spread $H = 333.8$, $p = 5.1 \times 10^{-60}$; tipward velocity $H = 461.5$, $p = 1.2 \times 10^{-86}$; shootward velocity $H = 466.2$, $p = 1.3 \times 10^{-87}$). Different letters denote significant differences among treatments

(Dunn's post hoc test with Benjamini–Hochberg correction, $p < 0.05$). Replicate numbers: NLP20 = 14; FLG22 = 79; C8 = 17; PG3 = 44; cellobiose = 17; PEP1 = 30; eATP = 29; L-Glu = 28; RALF23 = 20; NaCl = 61; KCl = 25; $Ca(NO_3)_2$ = 27; $KH_2PO_4$ = 15; $KNO_3$ = 22; $NH_4NO_3$ = 23; $(NH_4)_2HPO_4$ = 32; IAA = 24; D-sorbitol = 21; cold (½ MS) = 33.

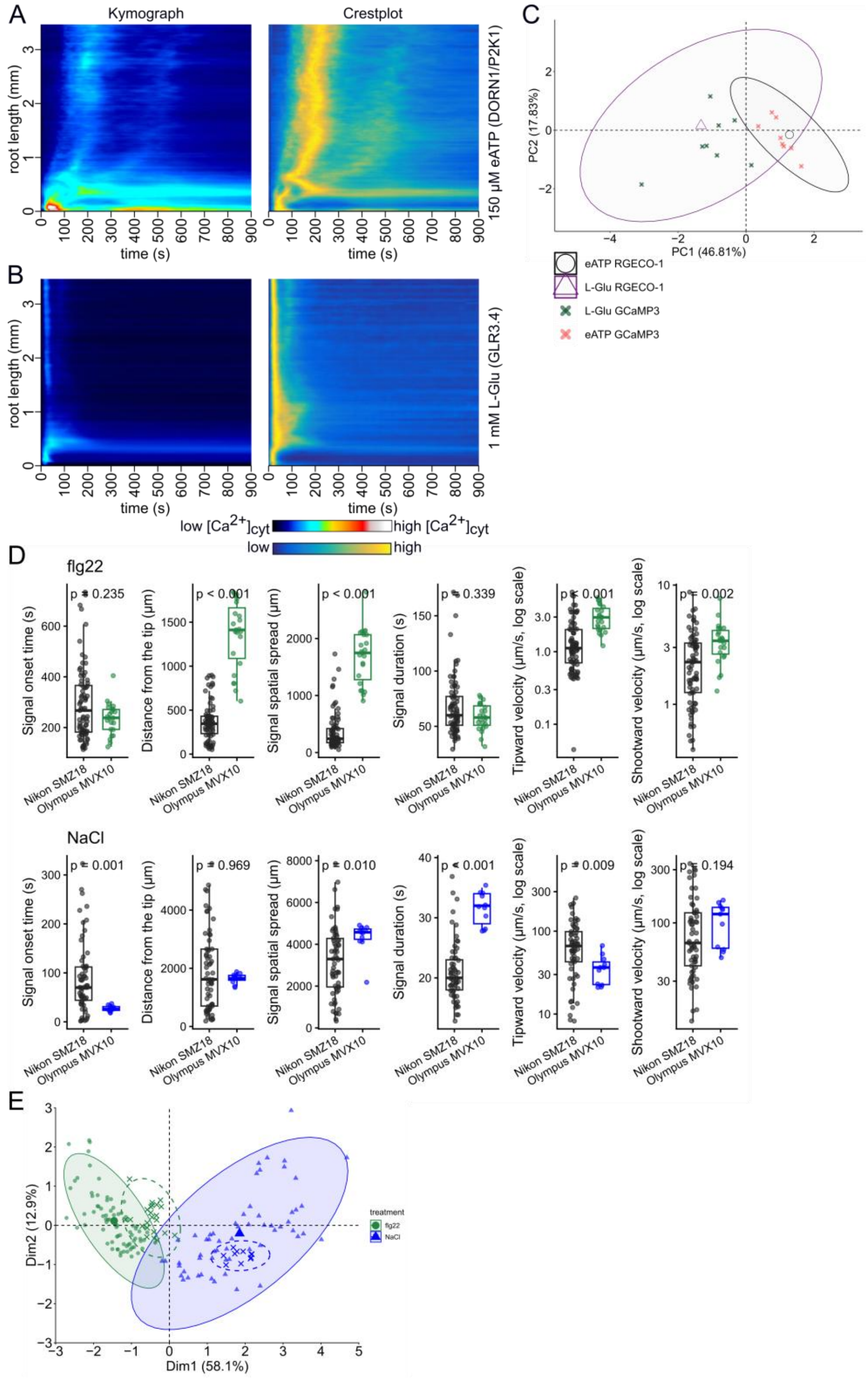


**Figure S5 Robustness of Crestline fingerprints across biosensors and imaging platforms.**

A) Representative kymograph (left) and corresponding crestplot (right) of cytosolic $Ca^{2+}$ dynamics in *Arabidopsis* roots expressing *35S::GCaMP3* following elicitation with 150 µM eATP (DORN1/P2K1-dependent; n = 8). B) As in (A), following elicitation with 1 mM L-glutamate (GLR-dependent; n = 8). In (A) and (B), the x-axis denotes time (1 pixel column = 1 s), and the y-axis denotes position along the root (root tip at the bottom). The royal LUT spans a fixed display range (min 100 to max 800, arbitrary units) applied identically to both kymographs, so intensities are directly comparable between them. The wave_tracking LUT spans the full 16-bit output range of the Crestline normalization (0 to 65,535), applied identically to both crestplots; values are normalized over time at each root position, so they represent relative signal within each position's time course rather than absolute $[Ca^{2+}]_{cyt}$. C) PCA of the six $Ca^{2+}$ fingerprint descriptors fitted on *UBQ10::R-GECO1* recordings for 150 µM eATP (n = 29) and 1 mM L-Glu (n = 28) from the $Ca^{2+}$tlas. *35S::GCaMP3* replicates (eATP n = 9, L-Glu n = 8; recorded on the Olympus MVX10) are projected as supplementary individuals and do not contribute to the PCA. Open symbols mark R-GECO1 group centroids; ellipses show 95 % confidence ellipses of the R-GECO1 groups. D) Fingerprint descriptors of wild-type *UBQ10::R-GECO1* roots treated with 10 µM FLG22 or 100 mM NaCl, recorded for the $Ca^{2+}$tlas (Nikon SMZ18; n = 79 and 61) or alongside the mutant experiments (Olympus MVX10; n = 23 and 11); Mann–Whitney tests, Benjamini–Hochberg-adjusted; the larger field of view of the Olympus allows signals to be followed over a longer root segment; velocities on a log axis for display only. E) PCA fitted on the $Ca^{2+}$tlas FLG22 and NaCl roots (filled symbols, solid 95 % normal ellipses), with Olympus roots projected as supplementary individuals (crosses, dashed ellipses).

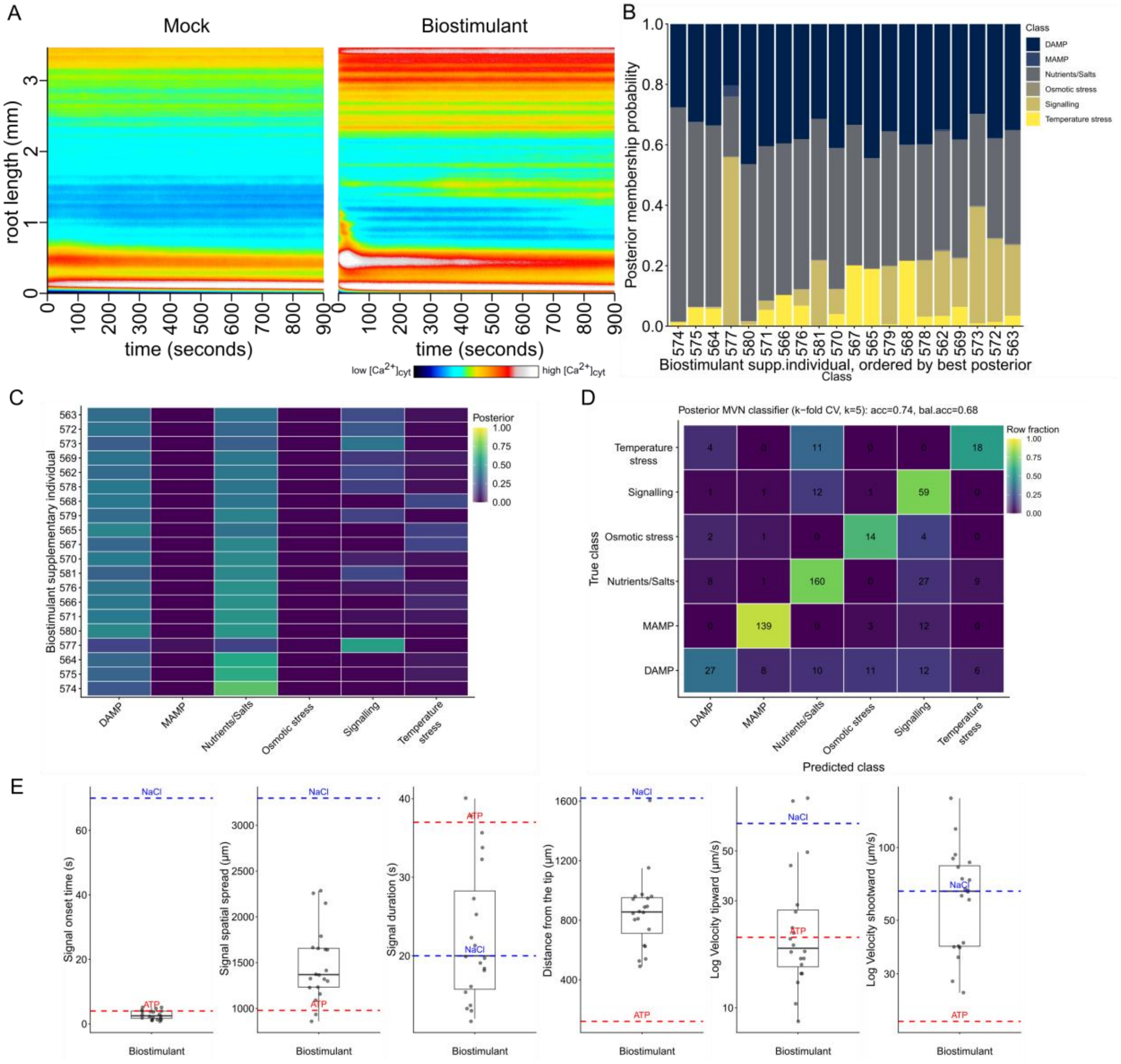


**Figure S6 How closely individual biostimulant replicates match each reference stimulus category, and how accurate this classification approach is.** A) Representative kymographs (royal LUT) of cytosolic $Ca^{2+}$ dynamics in wild-type Arabidopsis root tips expressing R-GECO1 following mock/biostimulant treatment. the x-axis denotes time (1 pixel column = 1 s), and the y-axis denotes position along the root (root tip at the bottom). The royal LUT spans a fixed display range (min 100 to 800, arbitrary units) applied identically to both kymographs, so intensities are directly comparable between them. B) Per-individual posterior membership of biostimulant supplementary individuals to the pre-defined elicitor classes, displayed as a stacked barplot (individuals ordered by maximum posterior probability). Under frequency-based priors, posterior support is split mainly between the Nutrients/Salts and DAMP classes, whereas MAMP and osmotic/temperature-stress classes contribute little. C) The same posterior probability matrix shown as a heatmap (rows = biostimulant data; columns = elicitor classes; colour indicates posterior probability), highlighting the consistency of class support across individuals and potential mixed-class assignments. D) Stratified 5-fold cross-validation of the posterior class-conditional multivariate normal (MVN) classifier on the active reference elicitor dataset. The confusion matrix reports row-normalized fractions (colour scale) with raw counts overlaid in each cell. Overall accuracy (acc = 0.74) and balanced accuracy (bal.acc = 0.68) are indicated in the title. Consistent with the diagonal structure, MAMP and nutrients/salts show high recall, while DAMP exhibits lower recall (greater overlap with other classes); corresponding class-wise recall values are provided in Table S2. E) Quantitative spatiotemporal descriptors of $Ca^{2+}$ fingerprints in *Arabidopsis* roots treated with the biostimulant. Boxplots show signal onset time, distance from the root tip, signal duration, spatial spread, and tipward/shootward velocities for individual roots. Dashed lines indicate the median response of representative Nutrients/Salts (NaCl) and DAMPs (eATP) treatments for comparison.

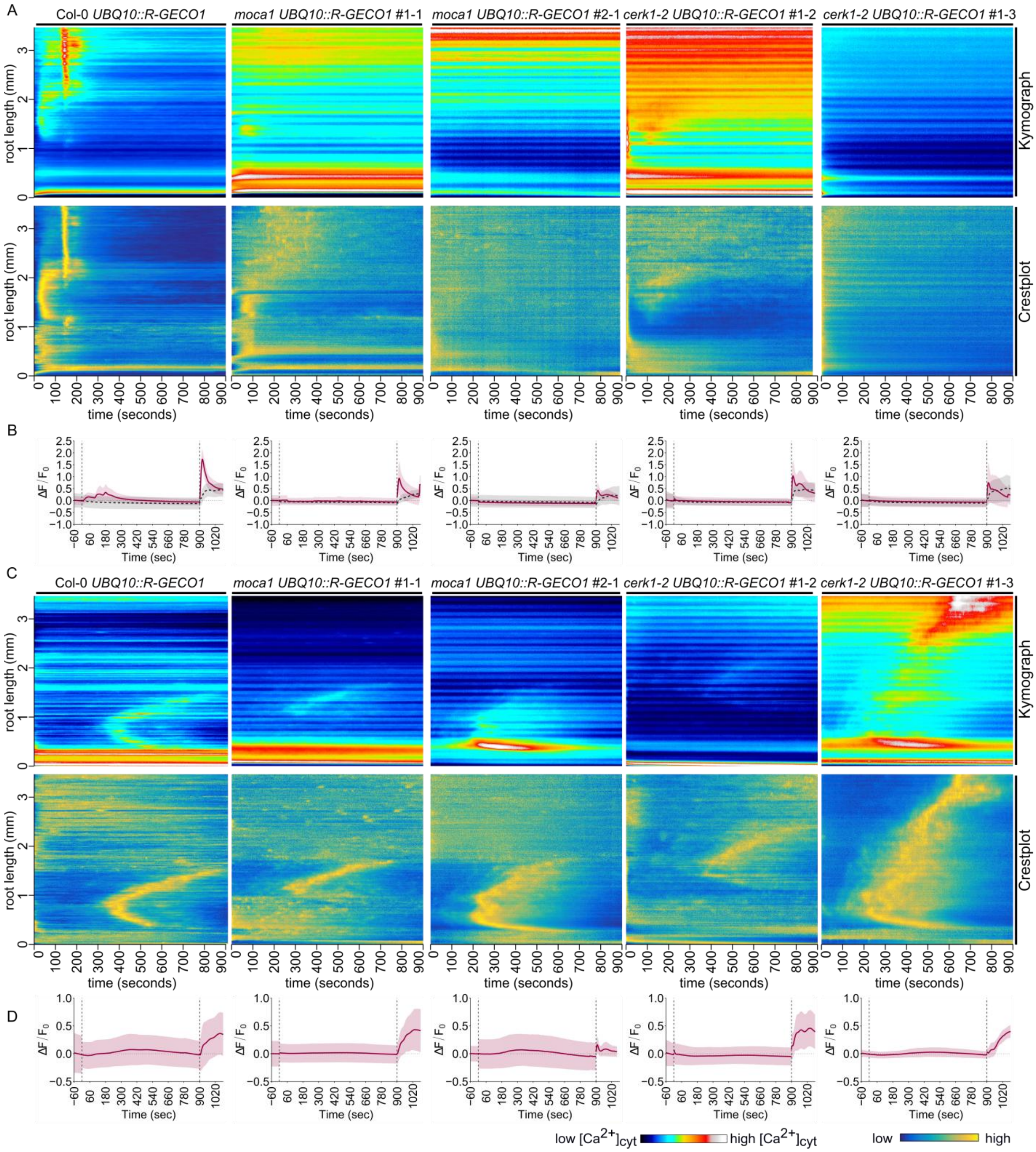


**Figure S7 Functional validation of R-GECO1 $Ca^{2+}$ reporter lines in mutant backgrounds.** Spatiotemporal $Ca^{2+}$ dynamics in *Arabidopsis* roots expressing *UBQ10::R-GECO1* in wild type (Col-0) and in two independent reporter lines for each mutant background (*moca1*, *cerk1-2*) were recorded following elicitation with 100 mM NaCl (a,b) or 10 µM FLG22 (c,d), followed by 150 μM eATP as a positive control. For each genotype, kymographs and corresponding crestplots are shown (a,c), together with $\Delta F/F_0$ traces (mean ± SD across replicates) aligned to stimulus application (vertical dashed lines) (b,d). $F_0$ is the mean of the 60 s pre-stimulus baseline; for the eATP phase, $F_0$ is the mean of the last 20 s before eATP application. (a,b) NaCl responses were used to verify the expected salt-induced $Ca^{2+}$ fingerprint phenotype in each background. The black dashed line in (b) indicates the mock treatment (½ MS medium). (c,d) FLG22, reported to act independently of the tested mutant backgrounds, was used as a second validation stimulus; all lines exhibited a clear FLG22-induced $Ca^{2+}$ response. In kymographs and crestplots, the x-axis denotes time (1 pixel column = 1 s), and the y-axis denotes position along the root (root tip at the bottom left). The royal LUT spans a fixed display range (min 100 to max 800, arbitrary units) applied identically to all kymographs, so intensities are directly comparable between them. The wave_tracking LUT spans the full 16-bit output range of the Crestline normalization (0 to 65,535), applied identically to all crestplots; values are normalized over time at each root position, so they represent relative signal within each position's time course rather than absolute $[Ca^{2+}]_{cyt}$.

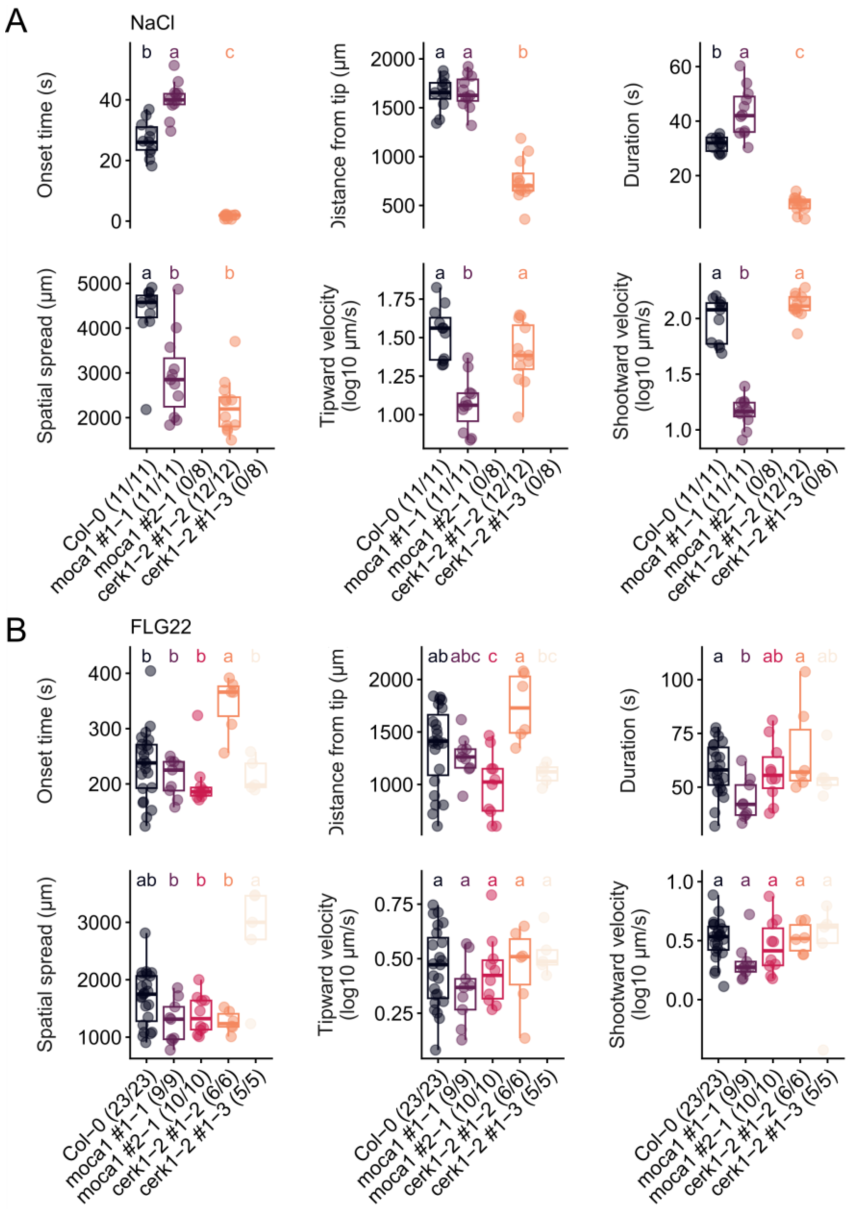


**Figure S8 Fingerprint descriptors of FLG22 and NaCl responses in the reporter lines.** (a) NaCl (100 mM) and (b) FLG22 (10 µM) responses of wild type (Col-0) and two independent UBQ10::R-GECO1 reporter lines each of moca1 (#1-1, #2-1) and cerk1-2 (#1-2, #1-3), from the experiments shown in Fig. S7. Numbers in parentheses give responding/total roots; non-responding roots are not plotted and were excluded from descriptor statistics. Kruskal–Wallis test followed by Dunn's post hoc test with Benjamini–Hochberg correction on responding roots; different letters denote significant differences ($p < 0.05$). NaCl response rates differed among lines (Fisher's exact test, $p < 0.0001$). Velocities are shown as $\log_{10}$ for display only; untransformed values were used for statistics.

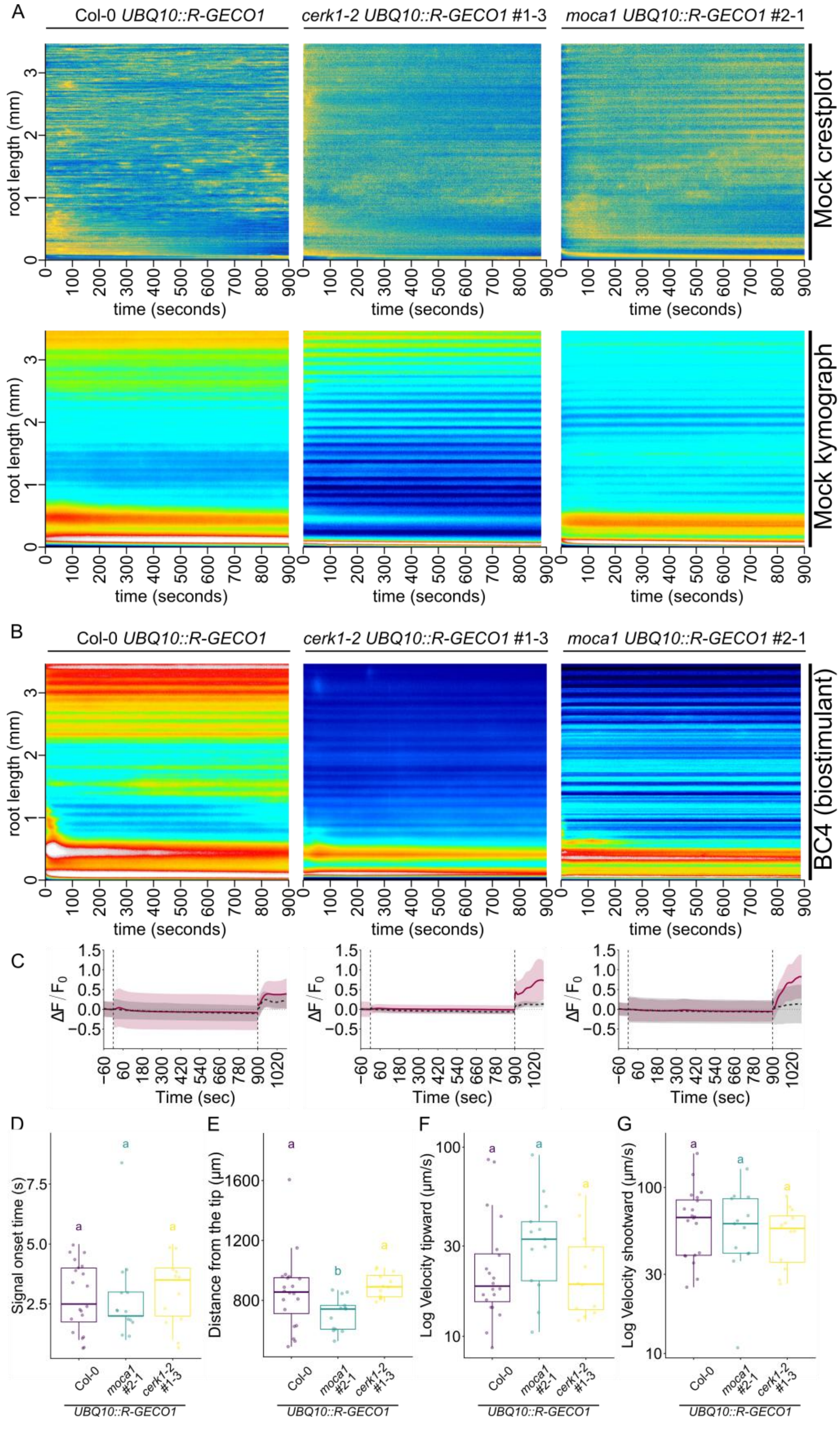


**Figure S9 Kymographs, intensity traces and fingerprint descriptors of BC4-evoked $Ca^{2+}$ signatures in wild type, *moca1* and *cerk1-2*.** A) Representative crestplots (top) and kymographs (bottom) of cytosolic $Ca^{2+}$ dynamics in *Arabidopsis* roots expressing *UBQ10::R-GECO1* in wild type (Col-0), *cerk1-2* (#1-3) and *moca1* (#2-1) backgrounds (left to right) following mock treatment (½ MS medium). B) Representative kymographs for the same genotypes following biostimulant treatment (BC4, 4 µg C $mL^{-1}$ in ½ MS medium). In (a) and (b), the x-axis denotes time (1 pixel column = 1 s), and the y-axis denotes position along the root (root tip at the bottom). The royal LUT spans a fixed display range (min 100 to max 800, arbitrary units) applied identically to all kymographs, so intensities are directly comparable between them. The wave_tracking LUT spans the full 16-bit output range of the Crestline normalization (0 to 65,535), applied identically to all crestplots; values are normalized over time at each root position, so they represent relative signal within each position's time course. C) Corresponding $\Delta F/F_0$ traces (mean ± SD) aligned to stimulus application (vertical dashed lines) for the sequential treatment ½ MS → mock or biostimulant → 150 µM eATP, providing the amplitude-based readout underlying the crestplots in Fig. 7A. The dashed black line indicates mock, and the magenta line indicates biostimulant treatment. $F_0$ is the mean of the 60 s pre-stimulus baseline; for the eATP phase, $F_0$ is the mean of the last 20 s before eATP application. D-G) Fingerprint descriptors not shown in Fig. 7: D) signal onset time, E) distance from the root tip at first detection, F) tipward and G) shootward velocity (log axis for display only). n = 20 (Col-0), 13 (*moca1* #2-1) and 12 (*cerk1-2* #1-3) roots. Letters denote groupings from Dunn's post hoc test with Benjamini–Hochberg correction ($p < 0.05$).

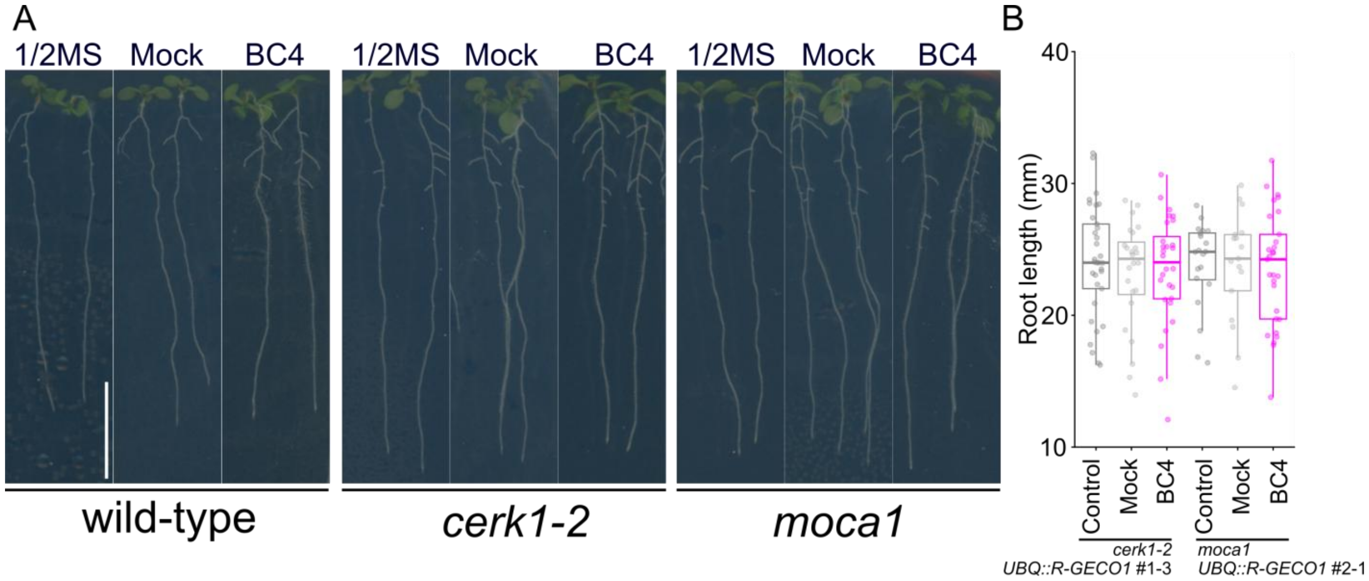


**Figure S10 Primary root growth under BC4 treatment.** A) Representative primary root growth phenotypes on ½ MS agar under control (½ MS), mock and BC4 (4 µg C $mL^{-1}$) treatment for wild type, cerk1-2 and moca1 (left to right). Scale bar = 10 mm. B) BC4 treatment did not significantly affect primary root length in the *cerk1-2* (#1-3) and *moca1* (#2-1) *UBQ10::R-GECO1* reporter lines, consistent with Fig. 7E. Two independent batches; n = 17–33 roots per condition.

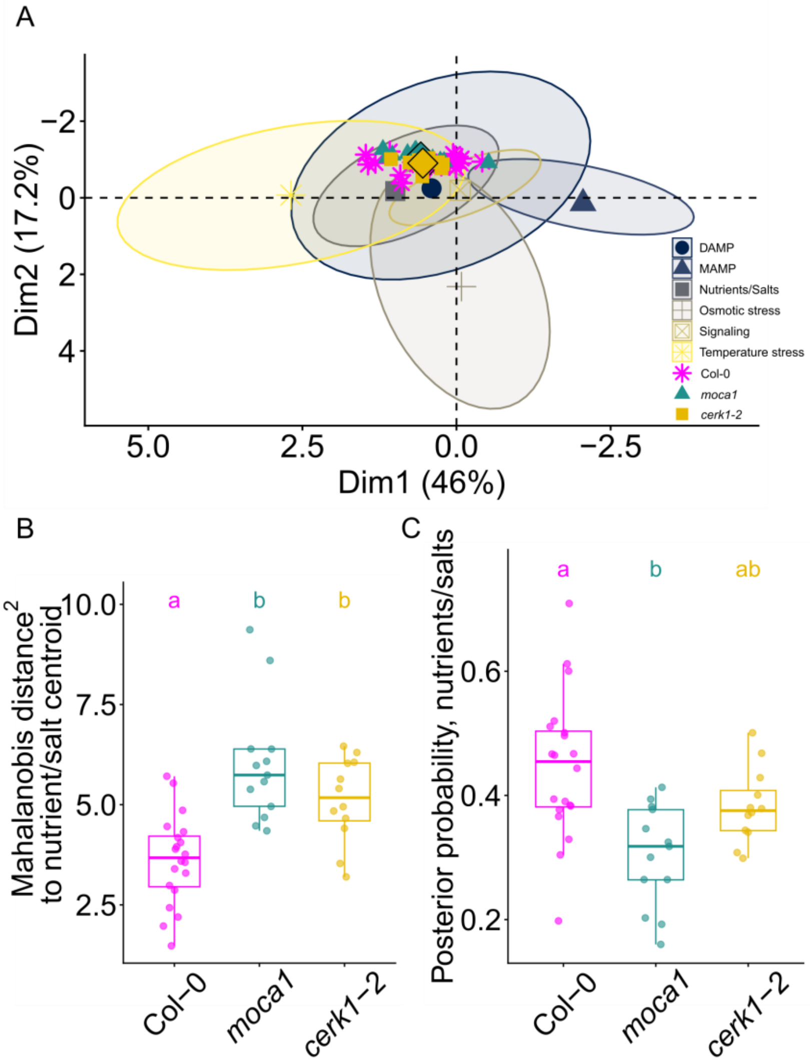


**Figure S11 BC4 responses of *moca1* and *cerk1-2* projected into the Ca²⁺tlas signaling space.** A) Reference stimulus classes (95% data ellipses, as in Fig. 6b) with BC4-treated roots of wild type (magenta asterisks), *moca1 UBQ10::R-GECO1* #2-1 (teal triangles) and *cerk1-2 UBQ10::R-GECO1* #1-3 (gold squares) projected as supplementary individuals; diamonds mark genotype centroids. B) Squared Mahalanobis distance of each BC4-treated root to the nutrient/salt class centroid in the first four principal components. C) Posterior probability of nutrient/salt class membership under frequency-based priors. In B) and C), Kruskal–Wallis test followed by Dunn's post hoc test with Benjamini–Hochberg correction; different letters denote significant differences ($p < 0.05$). n = 20 (wild type), 13 (*moca1*) and 12 (*cerk1-2*) roots. Data in Supplemental Datasets 3 and 4; statistics in Table S5.

**Table S1 | Elemental profile of the biostimulant (BC4) used in this study.** Elemental and physicochemical composition of the biostimulant stock (lot 6-23-25) as reported by fertilizer-grade analytical testing. Macro-nutrients are reported as % (w/w), trace metals as ppm, pH is measured according to EPA 150.1, and specific gravity according to ASTM D891-09. ND, not detected (below the method detection limit).

| Fertilizer Test | Method | Units | Results |
|---|---|---|---|
| Total Nitrogen (N) | AOAC 993.13 | % w/w | 0.51 |
| Available Phosphate ($P_2O_5$) | AOAC 3051A/6020 | % w/w | 1.38 |
| Soluble Potash ($K_2O$) | AOAC 3051A/6020 | % w/w | 0.17 |
| Calcium (Ca) | AOAC 3051A/6020 | % w/w | ND |
| Sodium (Na) | AOAC 3051A/6020 | % w/w | 0.028 |
| Sulfur (S) | AOAC 3051A/6020 | % w/w | ND |
| Magnesium (Mg) | AOAC 3051A/6020 | % w/w | ND |
| Iron (Fe) | AOAC 3051A/6020 | % w/w | 0.002 |
| Boron (B) | AOAC 3051A/6020 | % w/w | ND |
| Manganese (Mn) | AOAC 3051A/6020 | % w/w | ND |
| Copper (Cu) | AOAC 3051A/6020 | % w/w | ND |
| Zinc (Zn) | AOAC 3051A/6020 | % w/w | ND |
| Total Carbon (C) | AOAC 972.43 | % w/w | 0.060 |
| pH | EPA 150.1 | | 8.88 |
| Specific Gravity | ASTM D891-09 | | 1.016 |
| Arsenic (As) | AOAC 3051A/6020 | ppm | 0.19 |
| Aluminum (Al) | AOAC 3051A/6020 | ppm | 12.50 |
| Barium (Ba) | AOAC 3051A/6020 | ppm | ND |
| Cadmium (Cd) | AOAC 3051A/6020 | ppm | ND |
| Chromium (Cr) | AOAC 3051A/6020 | ppm | ND |
| Cobalt (Co) | AOAC 3051A/6020 | ppm | ND |
| Mercury (Hg) | AOAC 3051A/6020 | ppm | ND |
| Lead (Pb) | AOAC 3051A/6020 | ppm | ND |
| Nickel (Ni) | AOAC 3051A/6020 | ppm | 0.12 |
| Molybdenum (Mo) | AOAC 3051A/6020 | ppm | 0.13 |
| Selenium (Se) | AOAC 3051A/6020 | ppm | ND |

*Note (experimental working concentration): For imaging assays, the biostimulant was diluted to 4 µg C mL$^{-1}$ (4 mg L$^{-1}$ total C) – see Material and Methods section for the protocol. Based on the measured total carbon content (0.060% w/w) and specific gravity (1.016), this corresponds to an approximate 1:152 dilution of the stock. At this dilution, the formulation delivers approximately 1.3 mM total P (from 1.38% w/w $P_2O_5$, equivalent to ≈0.65 mM $P_2O_5$), ~2.4 mM total N (from 0.51% w/w N, not necessarily all ionic), ~0.24 mM $K^+$ (from 0.17% w/w $K_2O$), ~0.08 mM $Na^+$ (from 0.028% w/w Na) and ~3 µM Al (from 12.5 ppm), in addition to the ½ MS background medium.*

**Table S2 | Posterior MVN classifier performance by elicitor class (stratified 5-fold cross-validation).** Classification performance of the class-conditional multivariate normal (MVN) posterior model evaluated on the active reference elicitor dataset. For each elicitor class, the table reports the number of true samples (n) and the true-positive rate (recall), defined as the fraction of samples from that class correctly assigned to the same class (maximum posterior).

| Elicitor class | Number of samples | True-positive rate (recall) |
|---|---|---|
| MAMP | 154 | 0.903 |
| Signaling | 74 | 0.797 |
| Nutrients/Salts | 205 | 0.780 |
| Osmotic stress | 21 | 0.667 |
| Temperature stress | 33 | 0.545 |
| DAMP | 74 | 0.365 |

**Table S3 BC4 class assignment under frequency-based and uniform class priors.** For each reference class: prior probability under frequency-based priors, mean posterior probability across the 20 BC4 replicates under each prior, and number of replicates with maximum posterior in that class.

| Class | Prior (frequency-based) | Mean posterior, frequency priors | Mean posterior, uniform priors | BC4 replicates assigned, frequency priors | BC4 replicates assigned, uniform priors | Median percentile within class | Replicates within central 95% | Replicates nearest (Mahalanobis) | Mean kNN share |
|---|---|---|---|---|---|---|---|---|---|
| Nutrients/salts | 0.365 | 0.446 | 0.212 | 14 | 0 | 0.76 | 20 | 0 | 0.41 |
| DAMP | 0.132 | 0.360 | 0.461 | 4 | 16 | 0.07 | 20 | 19 | 0.42 |
| Signaling | 0.132 | 0.130 | 0.162 | 2 | 2 | 0.95 | 10 | 0 | 0.06 |
| Temperature stress | 0.059 | 0.060 | 0.157 | 0 | 2 | 0.80 | 17 | 1 | 0.09 |
| MAMP | 0.275 | 0.002 | 0.001 | 0 | 0 | 1.00 | 2 | 0 | 0.01 |
| Osmotic stress | 0.037 | 0.001 | 0.007 | 0 | 0 | 1.00 | 0 | 0 | 0.00 |

Legend: *Frequency-based priors are proportional to class size in the reference panel; uniform priors are 1/6 for each class. Mean posteriors are averaged over the 20 BC4 replicates; replicates are assigned to the class with maximum posterior probability. Nearest: number of BC4 replicates with minimum Mahalanobis distance to that class. Median percentile: for each replicate, the fraction of the class's own reference roots closer to the class centroid than the replicate; median over the 20 replicates (low values indicate a typical member). Within central 95%: number of replicates with a percentile below 0.95. Mean kNN share: fraction of each replicate's 23 nearest reference roots (Euclidean distance, first four PCs) belonging to the class, averaged over replicates.*

**Table S4 | Leave-one-elicitor-out validation.** For each held-out elicitor: true class, number of roots, and, under frequency-based and uniform class priors, the most frequently predicted class, the fraction of roots assigned to the true class, and the mean posterior probability of the true class. D-sorbitol and cold could not be evaluated because each is the only member of its class.

| **Held-out elicitor** | **True class** | **n roots** | **Predicted class (most frequent), frequency priors** | **Fraction of roots correct, frequency priors** | **Mean posterior, true class, frequency priors** | **Predicted class (most frequent), uniform priors** | **Fraction of roots correct, uniform priors** | **Mean posterior, true class, uniform priors** |
|---|---|---|---|---|---|---|---|---|
| nlp20 | MAMP | 14 | MAMP | 1.00 | 1.00 | MAMP | 1.00 | 1.00 |
| flg22 | MAMP | 79 | MAMP | 1.00 | 0.99 | MAMP | 1.00 | 0.99 |
| C8 | MAMP | 17 | MAMP | 0.59 | 0.56 | MAMP | 0.59 | 0.54 |
| PG3 | MAMP | 44 | MAMP | 0.50 | 0.48 | Signaling | 0.48 | 0.45 |
| Cellobiose | DAMP | 17 | Osmotic stress | 0.00 | 0.01 | Osmotic stress | 0.00 | 0.01 |
| eATP | DAMP | 29 | Signaling | 0.00 | 0.08 | Signaling | 0.14 | 0.16 |
| L-Glu | DAMP | 28 | Nutrients/ salts | 0.00 | 0.06 | Temperature stress | 0.11 | 0.10 |
| PEP1 | Signaling | 30 | Signaling | 0.70 | 0.48 | Signaling | 0.77 | 0.65 |
| RALF23 | Signaling | 20 | Signaling | 0.85 | 0.60 | Signaling | 1.00 | 0.75 |
| IAA | Signaling | 24 | Nutrients/ salts | 0.17 | 0.16 | DAMP | 0.21 | 0.19 |
| NaCl | Nutrients/ salts | 61 | DAMP | 0.15 | 0.16 | DAMP | 0.11 | 0.11 |
| KCl | Nutrients/ salts | 25 | Nutrients/ salts | 0.80 | 0.66 | Nutrients/ salts | 0.56 | 0.47 |

| Held-out elicitor | True class | n roots | Predicted class (most frequent), frequency priors | Fraction of roots correct, frequency priors | Mean posterior, true class, frequency priors | Predicted class (most frequent), uniform priors | Fraction of roots correct, uniform priors | Mean posterior, true class, uniform priors |
|---|---|---|---|---|---|---|---|---|
| $Ca(NO_3)_2$ | Nutrients/ salts | 27 | Nutrients/ salts | 0.74 | 0.66 | Nutrients/ salts | 0.70 | 0.47 |
| $KH_2PO_4$ | Nutrients/ salts | 15 | Nutrients/ salts | 0.80 | 0.70 | Nutrients/ salts | 0.60 | 0.54 |
| $KNO_3$ | Nutrients/ salts | 22 | Nutrients/ salts | 0.95 | 0.75 | Nutrients/ salts | 0.86 | 0.56 |
| $NH_4NO_3$ | Nutrients/ salts | 23 | Nutrients/ salts | 0.83 | 0.73 | Nutrients/ salts | 0.70 | 0.60 |
| $(NH_4)_2HPO_4$ | Nutrients/ salts | 32 | Nutrients/ salts | 0.78 | 0.72 | Nutrients/ salts | 0.69 | 0.59 |
| D-sorbitol | Osmotic stress | 21 | not evaluable | – | – | not evaluable | – | – |
| Cold | Temperature stress | 33 | not evaluable | – | – | not evaluable | – | – |

*Legend: All replicates of one elicitor were withheld, the PCA and class models were refit on the remaining elicitors with frequency-based or uniform priors, and the withheld elicitor was projected and assigned to the class predicted for most of its replicates. Twelve of 17 evaluable elicitors were assigned to their correct class under frequency-based priors (root-level accuracy 0.58), and 11 of 17 under uniform priors (root-level accuracy 0.56). D-sorbitol and cold could not be evaluated because each is the only member of its class.*

**Table S5 | Position of BC4 responses in the Ca²⁺tlas signaling space by genotype.** BC4-treated roots of wild type (n = 20), *moca1* #2-1 (n = 13) and *cerk1-2* #1-3 (n = 12) projected into the reference PCA (first four PCs). (a) Mean posterior class membership (frequency-based / uniform priors). (b) Median squared Mahalanobis distance to each class centroid.

**(a) Mean posterior class membership, frequency-based / uniform priors**

| Genotype | DAMP | MAMP | Nutrients/salts | Osmotic stress | Signaling | Temperature stress |
|---|---|---|---|---|---|---|
| Col-0 | 0.36 / 0.46 | 0.00 / 0.00 | 0.45 / 0.21 | 0.00 / 0.01 | 0.13 / 0.16 | 0.06 / 0.16 |
| *moca1* | 0.53 / 0.57 | 0.01 / 0.00 | 0.30 / 0.12 | 0.00 / 0.00 | 0.07 / 0.08 | 0.09 / 0.21 |
| *cerk1-2* | 0.46 / 0.53 | 0.00 / 0.00 | 0.38 / 0.16 | 0.00 / 0.01 | 0.08 / 0.10 | 0.08 / 0.20 |

**(b) Median squared Mahalanobis distance to each class centroid**

| Genotype | DAMP | MAMP | Nutrients/salts | Osmotic stress | Signaling | Temperature stress |
|---|---|---|---|---|---|---|
| Col-0 | 0.96 | 40.87 | 3.68 | 11.81 | 9.71 | 4.72 |
| *moca1* | 1.69 | 59.62 | 5.74 | 20.03 | 20.47 | 4.95 |
| *cerk1-2* | 2.06 | 38.11 | 5.17 | 13.48 | 11.40 | 4.30 |
